# A Unified Framework for Selecting and Evaluating Cell-Type-Specific Gene Co-expressions in Single-Cell Data

**DOI:** 10.1101/2024.11.08.622700

**Authors:** Xinning Shan, Yingxin Lin, Hongyu Zhao

## Abstract

Cell-type-specific gene co-expression networks are widely used to characterize gene relationships. Although many methods have been developed to infer such co-expression networks from single-cell data, the lack of consideration of false positive control in many evaluations and downstream analyses may lead to incorrect conclusions because higher reproducibility, higher functional coherence, and a larger overlap with known biological networks may not imply better performance if the false positives are not well controlled. In this study, we systematically compared two distinct criteria for selecting correlated gene pairs from single-cell data, p-value versus correlation strength. We found that the use of p-values instead of correlation strength is more robust for both selecting meaningful gene pairs and for the fair benchmarking of co-expression estimation methods. To make this approach universally applicable, we extended and validated a simulation method that can efficiently and reliably generate empirical p-values for co-expression estimation methods that do not have corresponding or well-controlled p-values. Furthermore, we demonstrated that a fair comparison of the estimation methods requires adjusting for the varying number of gene pairs they identified and accounting for the inherent expression-level biases within ground truth biological networks. Our study provides a practical guide for researchers to select reliable correlated gene pairs for downstream study and establishes a more rigorous standard for the evaluation and comparison of gene co-expression network estimation methods.

## 1 Background

With the development of single-cell technology and the potential of inferring cell-type-specific gene coexpression networks from these data to understand relationships among genes, many methods have been developed for estimating such networks through single-cell RNA-sequencing (scRNA-seq) data. These methods typically result in a dense network with a value for each possible gene pair. Therefore, the accurate identification of co-expressed gene pairs from the initial network is a critical step for downstream analyses and method benchmarking. However, this post-selection step introduces two key challenges. First, there is no consensus approach for selecting truly co-expressed gene pairs. Second, there is no standard for the fair comparison of the gene co-expression estimation methods.

There are two commonly used approaches for selecting correlated gene pairs in real data analysis, p-value-based and correlation-strength-based. The correlation-strength-based approach selects correlated gene pairs based on the estimated correlation strength, and is more commonly used because of its convenience [1–8]. However, the correlation-strength-based approach does not consider false positive control. Although a p-value-based approach is theoretically more robust, a formal comparison to quantify the practical difference between these two approaches is lacking in the context of cell-type-specific single-cell analysis to the best of our knowledge. Furthermore, a rigorous p-value-based approach is hampered by the fact that many methods do not provide p-values or the p-value is not well-controlled [9]. These issues are further complicated during method evaluation. Standard benchmarks are often biased not only because different methods identify widely varying numbers of correlated gene pairs, but also because the biological databases used as ground truth can have their own expression biases, making direct comparisons unfair.

To address these critical gaps, we propose a unified, p-value-driven framework for post-selecting and evaluating cell-type-specific gene co-expressions. We first establish our primary finding that p-value-based thresholding is superior to correlation-strength-based thresholding using one exemplar of the co-expression network estimation method, which uniquely provides both estimates and well-calibrated p-values (Section 2.1). To generalize this finding, we address the lack of well-calibrated p-values in other co-expression estimation methods by generating empirical p-values. To do this robustly, we first conduct a benchmark of simulation methods and identify scSimu as the optimal choice for this task (Section 2.2). Next, using scSimu to generate well-calibrated empirical p-values for all methods, we generalize our initial finding in Section 2.1 (Section 2.3). Finally, we extend this framework to enable fair method comparison, introducing a necessary adjustment that accounts for method-specific differences in statistical power (Section 2.4). Our work provides both a practical workflow for researchers and a more rigorous standard for the benchmarking of gene co-expression estimation methods.

## 2 Results

### 2.1 A lack of type I error control in identifying correlated gene pairs can lead to misleading results

In this section, we demonstrate the critical need for using p-values to control false positives in the evaluation of gene co-expression network estimation methods. We focused our analysis on CS-CORE, a method that provides well-calibrated p-values for cell-type-specific gene co-expressions [9]. Because we do not know the ground truth of correlated gene pairs in real data, we used simulations that mimicked real data to better understand the impact of appropriate control of false positives (see Methods for details). For the evaluation measures, we focused on two most commonly used ones, reproducibility [1, 4, 5, 10] and overlap with known biological networks (STRING [11] and Reactome [12]) [1, 2, 4, 6, 7, 13].

We used these simulations to directly compare the performance of the correlation-strength and p-value-based approaches. In the correlation-strength-based approach, selecting a larger number of top pairs leads to higher reproducibility and more overlaps with biological networks (Figures 1A and 1F). However, it may include more false positives. As shown in Figures 1A and 1F, an increasing number of reproducible pairs and a higher number of overlaps with STRING are both associated with a lower precision. We also observe higher reproducibility and more overlaps when the p-value-based approach with a less stringent cut-off is used (Figures 1B and 1G), but there is a less decline in precision (Figures 1B and 1G).

**Fig. 1.**
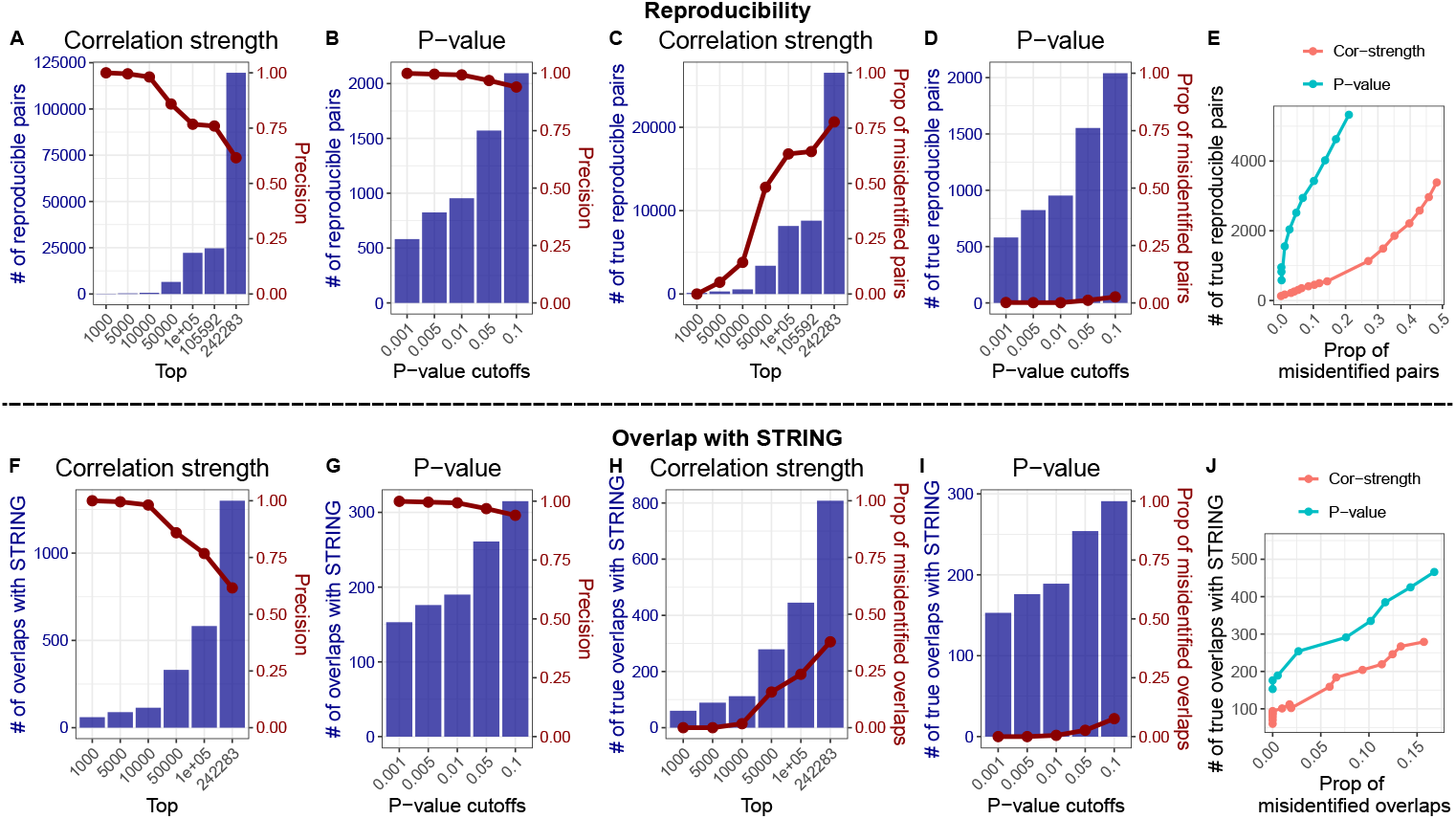
Comparison of correlation-strength and p-value-based approaches for selecting co-expressed gene pairs. Using results from CS-CORE, the performance of the correlation-strength-based and the p-value-based approach was evaluated by reproducibility (A-E) and overlap with the STRING database (F-J). **A-B**. Total number of reproducible pairs (blue bars, left y-axis) and precision (red line, right y-axis) for the correlation-strength (A) and p-value (B) approaches. The x-axis represents the number of top pairs selected by correlation strength or the p-value cutoff. (Precision in PNAS data can be found in Figure S1) **C-D**. Number of true reproducible pairs (blue bars, left y-axis) and the proportion of misidentified pairs, 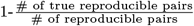 (red line, right y-axis). **E**. Direct comparison of the number of true reproducible pairs versus the proportion of misidentified pairs. **F-G**. Total number of overlaps with the STRING and precision for the correlation-strength (F) and p-value (G) approaches. **H-I**. Number of true overlaps with STRING and the proportion of misidentified overlaps for the correlation-strength (H) and p-value (I) approaches. **J**. Direct comparison of the number of true overlaps versus the proportion of misidentified overlaps.

To understand the influence of these false positives in the evaluation process, we investigated the proportion of misidentified reproducible pairs (see Methods for details). As expected, there is an increased proportion of misidentification with a growing number of gene pairs selected (Figure 1C). We find that the proportion of misidentified reproducible pairs based on p-value-based approach is much lower than that based on the correlation-strength-based approach (Figure 1D). The results are similar to those comparing overlaps with STRING (Figures 1H-I) and overlaps with Reactome (Figure S2). Even though the correlation-strength-based approach has a much higher number of reproducible pairs and overlaps with known biological networks, the large discrepancy in precision and proportion of misidentification at different thresholds necessitates more careful threshold selection. Additionally, variations in the density and magnitude of the correlation matrix across datasets further complicate threshold selection, making applying the same threshold to different datasets questionable.

As shown in Figure 1, there is a potential trade-off between the proportion of false identification and the number of true reproducible pairs or true overlaps with known biological networks, i.e., a lower proportion of false identification may correspond to a lower number of true reproducible pairs or true overlaps. To better quantify this, we compared the number of true reproducible pairs identified by the correlation-strength-based approach and the p-value-based approach, ensuring that their proportions of false identifications are the same in Figure 1E, which further supports that selecting correlated gene pairs based on p-value is better than that based on correlation strength. Similarly, we observe a higher number of true overlaps with known biological networks when using the p-value-based approach (Figures 1J and S2E).

### 2.2 A validated simulation to enable p-value selection across all methods

While our results demonstrated the superiority of using p-value-based approach, its practical application is often limited as many co-expression estimation methods do not provide p-values. Among those that do, p-values are often derived from theoretical null distributions, such as the Student’s t-distribution, which may not be suitable for sparse and complex scRNA-seq data. This is further supported by Su et al. [9] which examined the p-value distribution of many co-expression estimation methods on permuted single-cell RNA count data. It was found that only Normalisr [14], Pearson correlation on the sctransform normalized data [15] and CS-CORE have well-calibrated type I errors [9].

Even though Pearson correlation on sctransform normalized data was shown to have well-calibrated type I errors, the p-values are derived from a theoretical Student’s t-distribution. As shown by 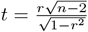, this p-value only depends on the correlation coefficient (*r*) and the number of single cells used (*n*). This means that two gene pairs with the same r and n will yield identical p-values, regardless of other inherent differences in their expression distributions, such as mean and variance (Figures S3A and S3B). There are similar concerns for the Spearman correlation. Therefore, a data-driven empirical p-value, which can account for these specific data characteristics, is necessary to provide a more robust and appropriate measure of statistical significance. To address this need, we employed a simulation-based framework to generate well-controlled empirical p-values, thereby enabling a rigorous and significance-based selection for any coexpression estimation methods.

Many methods have been developed for simulating count data in the context of scRNA-seq. Crowell et al. [16] found that ZINB-WaVE [17] and scDesign2 [18] were the top two methods for simulating data containing cells from a single batch and cluster. In another benchmark paper, ZINB-WaVE and SPARSim [19] were found to exhibit greater similarity to the original data in terms of both data properties and biological signals [20]. More recently, scDesign2 has been further developed into scDesign3 [21]. Even though these methods can closely mimic real data, some limitations exists. ZINB-WaVE [17] does not consider the correlation structure, whereas scDesign2 [18] and scDesign3 [21] are computationally demanding requiring several hours for a single simulation (Figures 2C and 2D). SPARSim’s [19] assumption of a fixed sequencing depth per cell is a key limitation, as this does not align with the stochastic nature of sequencing in real datasets. Therefore, we introduce scSimu (Single-Cell SIMUlation), an improved and extended version of the simulation method used in our previous publication [9] (see Methods for details), which serves as an accurate and efficient tool for evaluating co-expression networks by simulating realistic scRNA-seq count data under the null hypothesis.

**Fig. 2.**
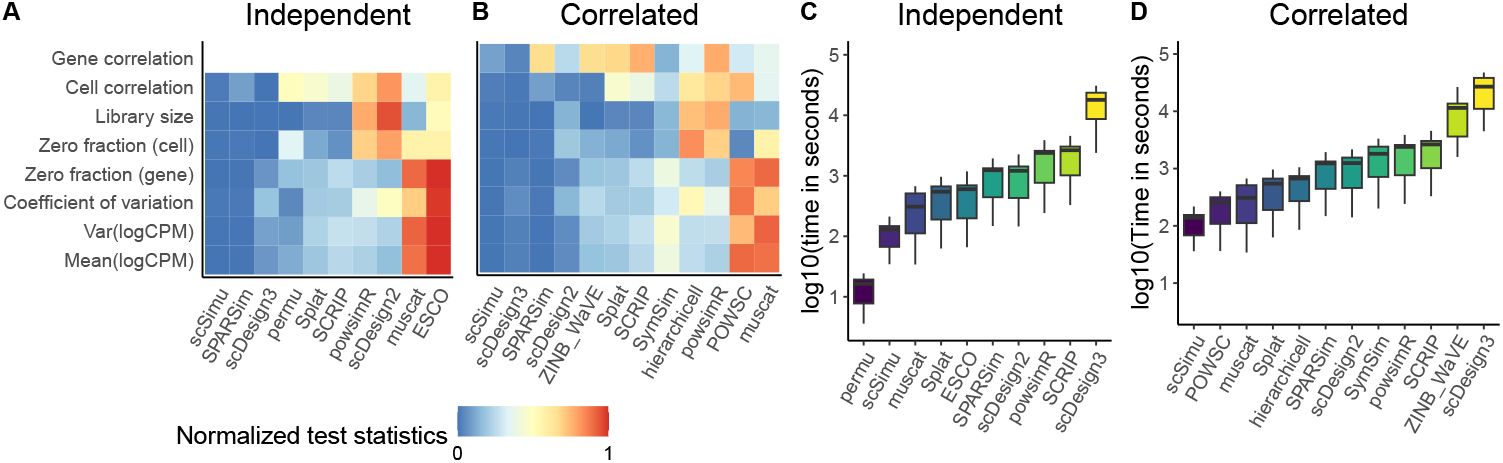
Benchmark of the simulation methods. **A and B**. The x-axis is the simulation method. The y-axis is the evaluation metric. The color represents the normalized test statistics of the combination of the KDE test and the KS test. Methods on the x-axis are ordered by the average normalized test statistics. **A**. Performance of simulating independent genes. **B**. Performance of simulating correlated genes. **C and D**. Boxplots of the computational time for simulating independent and correlated genes respectively, across 10 datasets. The y-axis represents the computational time, measured in seconds and displayed on a log10 scale. We specified the marginal distribution as negative binomial to improve the speed in scDesign2.

To validate the accuracy of scSimu in mimicking real data, we benchmarked its ability to simulate both independent and correlated genes against a total of 13 other simulation methods (Table S1; see Methods for details). As a baseline check, we first confirmed that all simulation methods in the independent comparison successfully generated a null dataset of independent genes (Table S2 and Figure S4). As shown in Figures 2A and 2B, scSimu, SPARSim, and scDesign3 achieve the greatest similarity with real data for both tasks. This is in line with the observation of Song et al. [21] that scDesign3 had better performance and that of Cao et al.

[20] where SPARSim was the second best method. However, ZINB-WaVE, the best method in two benchmark papers [16, 20], is not among the top three methods in our comparisons. Our benchmark exclusively evaluates methods on data from droplet-based technologies, as their high throughput and cost-effectiveness make them widely used in the field. The model assumption for scRNA-seq data differs across sequencing platforms. For example, a zero-inflated model is used for the Smart-Seq data [22]. In the two benchmark papers [16, 20], different types of scRNA-seq data were considered, likely contributing to conclusions that were inconsistent with ours. Although scSimu, SPARSim, and scDesign3 show similar performance, SPARSim’s generating mechanism is unrealistic and scDesign3’s extensive computational time makes it impractical for simulating null distributions (Figures 2C and 2D). Separated results for the KDE and KS tests (Figure S5) confirm these findings.

Based on its balance of accuracy and computational efficiency, we selected scSimu as the null data generator for our p-value framework. To provide a further validation for this purpose, we examined the distribution of empirical p-values of independent gene pairs from a scSimu generated real data simulation. As shown in Figure 3, the resulting p-value distribution for independent gene pairs almost aligns with a Uniform(0,1) distribution. Furthermore, this holds true for all seven gene co-expression estimation methods we considered, regardless of whether they originally provided p-values. This demonstrates that scSimu provides effective and consistent control over type I error, making it a reliable tool for generating the empirical p-values needed to test for significant co-expression.

**Fig. 3.**
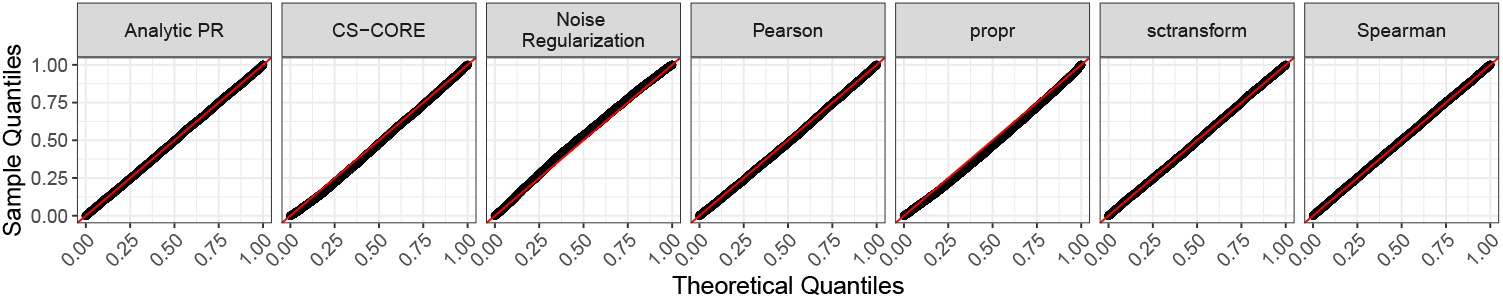
Q-Q plots of empirical p-values for independent gene pairs. Empirical p-values were calculated according to the methods outlined in Section 4.4.

### 2.3 P-value-based approach consistently outperforms and avoids benchmark distortions

To determine if the superiority of the p-value approach is a general principle, we extended our comparison to several other co-expression estimation methods. In Section 2.1, we showed that a p-value-based approach is better than the correlation-strength-based approach under CS-CORE. In this section, we extended this argument to a more general case by comparing the correlation-strength-based approach and the p-value-based approach using empirical p-values for six additional methods, including Noise Regularization [6], Pearson correlation on log-transformed data (Pearson), Spearman correlation on log-transformed data (Spearman), Pearson correlation on the analytic Pearson residuals (Analytic PR) [23], and Pearson correlation on the normalized data obtained via sctransform (sctransform). We also included CS-CORE using empirical p-values to further test if the previously drawn conclusion is specific to CS-CORE provided p-values.

The results are consistent with the findings from Section 2.1. For every method tested, the correlation-strength-based approach identifies a larger number of reproducible pairs and overlaps with biological networks, but at the cost of significantly lower precision and a higher proportion of misidentified pairs (Figures 4, 5, 6, S7, S8, and S9). In addition, the proportion of misidentification is especially high for certain gene co-expression estimation methods (Figure 4D). This finding indicates that some gene co-expression estimation methods, when using a correlation-strength-based approach, may seem artificially more effective due to not accounting for false positives. We estimated empirical p-values through scSimu and evaluated the p-value-based approach. Conversely, the p-value-based approach consistently maintains a lower proportion of false identifications, even when the p-value cutoff is relaxed to select more pairs (Figures 4, 5, 6, S7, S8, and S9). This confirms that the advantage of using statistical significance to control for false positives is a robust and generalizable finding, not specific to a single co-expression estimation method.

**Fig. 4.**
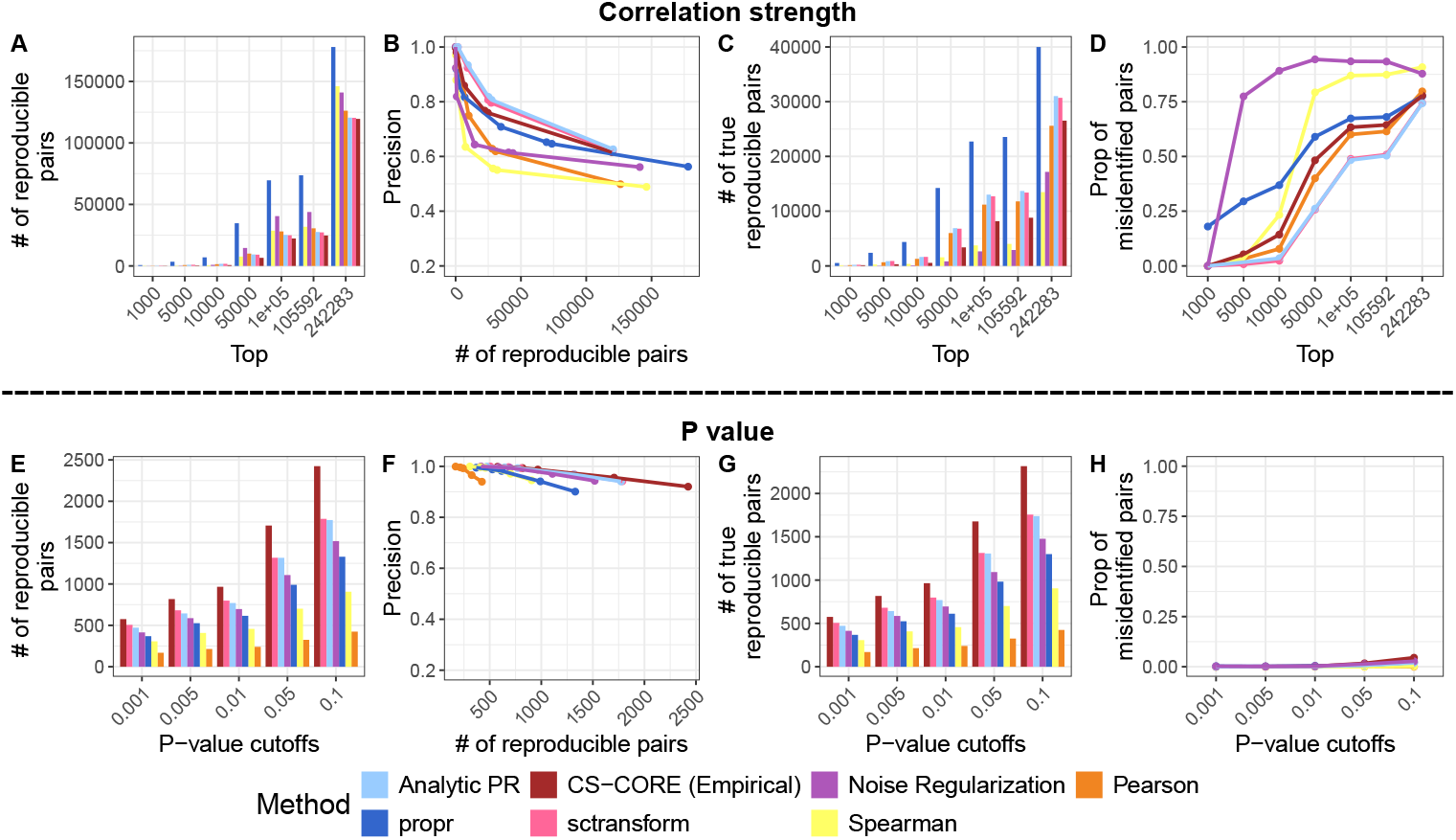
Performance in terms of reproducibility. Different estimation methods are represented with different colors. We use the term CS-CORE (Empirical) to denote the CS-CORE when its p-values is from our scSimu framework. Correlated gene pairs were identified via the correlation-strength-based approach (A-D), and p-value-based approach (E-H). **A and E**. The number of identified reproducible pairs at different cutoffs. **B and F**. Relationship between precision in ROSMAP data (precision in PNAS data can be found in Figure S6) and the number of reproducible pairs. Points in B and F correspond to different cutoffs for the correlation strength and p-value, respectively. **C and G**. The number of true reproducible pairs at different cutoffs. **D and H**. The proportion of misidentified reproducible pairs when gene pairs were selected by their estimated correlations (D) or by the empirical p-values (H).

**Fig. 5.**
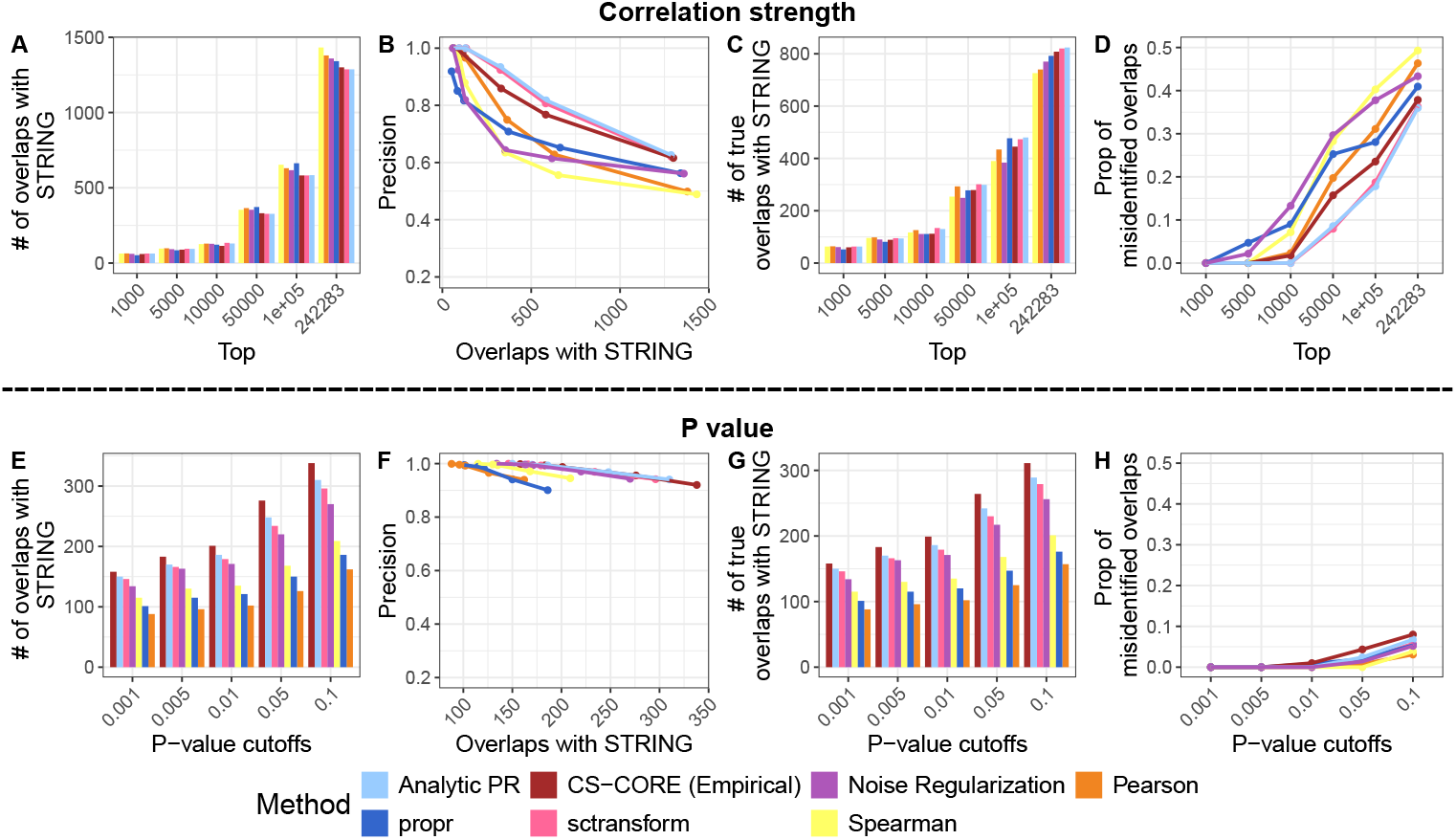
Performance in terms of overlaps with STRING. **A and E**. The number of overlaps with STRING at different cutoffs. **B and F**. Relationship between precision and the number of overlaps with STRING. **C and G**. The number of true overlaps with STRING at different cutoffs. **D and H**. The proportion of misidentified overlaps when gene pairs were selected by their estimated correlations (D) or by the empirical p-values (H).

**Fig. 6.**
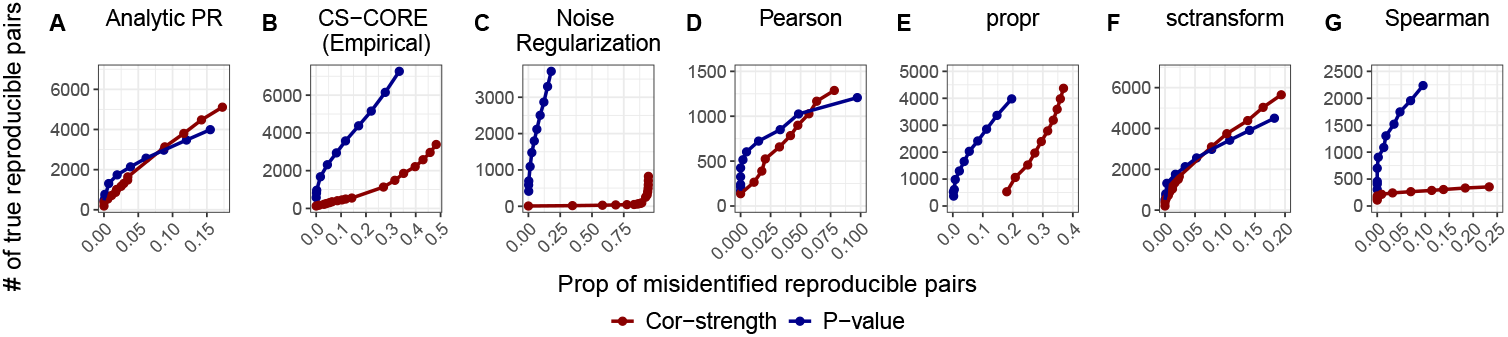
Comparison of correlated gene pair selection approaches in terms of reproducibility. The performance of selecting gene pairs by correlation strength (dark red) versus p-value (dark blue) was evaluated for seven different co-expression estimation methods. Each panel plots the number of true reproducible pairs (y-axis) against the proportion of misidentified reproducible pairs (x-axis) at varying selection thresholds.

Without considering the p-value in selecting correlated gene pairs can also affect the benchmark results across gene co-expression estimation methods. The ranks of estimation methods at the same cutoff vary substantially in the number of reproducible pairs and the number of true reproducible pairs when we use the correlation-strength-based approach (Figures 4A and 4C), i.e., a method that exhibits the highest reproducibility may not be the best method since many of the reproducible pairs may be false positives. On the contrary, the ranks are rather consistent when we use the p-value-based approach (Figures 4E and 4G). We observe the same pattern through overlaps with known biological networks (Figures 5 and S7). Our observation also explains the inconsistent results between the benchmark study of gene association estimation methods conducted by Skinnider et al. [1] which found propr to be superior, and the study conducted by Su et al. [9] where propr did not perform well, due to the lack of controls of false positives. Our study highlights the importance of considering statistical significance in gene pair selection. By selecting correlated gene pairs based on the p-value, as we demonstrated above, more reliable evaluation results can be achieved.

We further compared the two approaches for selecting correlated gene pairs using real data, focusing on oligodendrocytes and excitatory neurons in the ROSMAP data. To perform the comparison, we calculated the number of shared Gene Ontology (GO) terms for gene pairs identified exclusively by either the p-value-based approach or the correlation-strength-based approach. A higher number of shared GO terms indicates a stronger likelihood that two genes are correlated. Overall, gene pairs uniquely identified by the p-value-based approach exhibit a higher number of shared GO terms, suggesting that the p-value-based approach identifies more reliable correlated gene pairs (Figures 7 and S10).

**Fig. 7.**
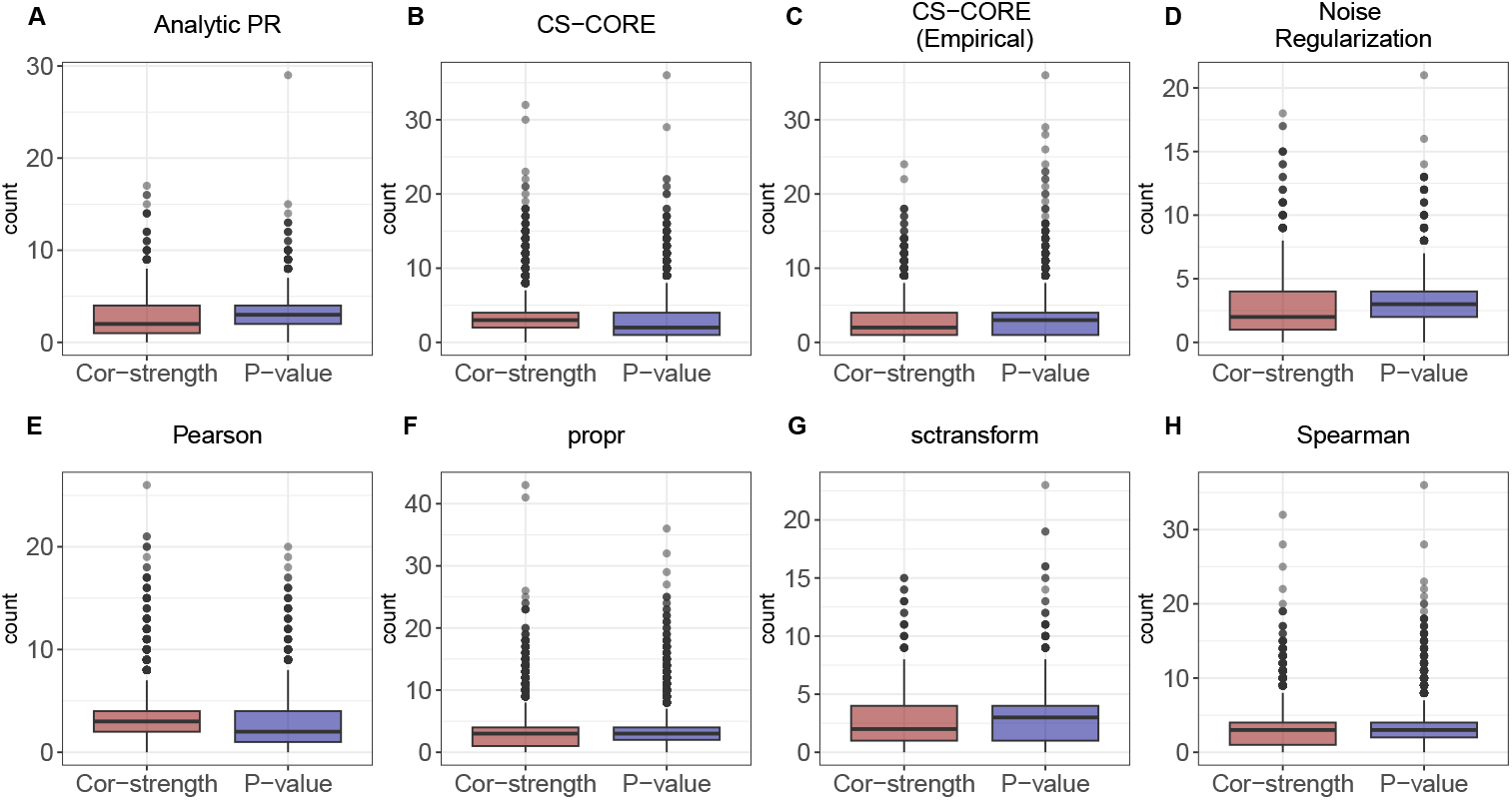
Comparison of shared GO terms for gene pairs identified exclusively by p-value or correlation-strength approaches. For each estimation method, we identified two sets of gene pairs in excitatory neurons, those found only by the p-value-based approach and those found only by the correlation-strength-based approach. To ensure a fair comparison, the threshold for the correlation-strength-based approach was set to yield the same number of gene pairs as the p-value-based approach at a 0.05 significance level (BH adjusted). The boxplots display the distribution of the number of shared GO terms for the gene pairs in each exclusive set.

### 2.4 Consideration of the overlap by chance and bias in the biological networks in the evaluation process

The p-value-based approach will cause a large discrepancy in the number of identified correlated gene pairs between different estimation methods. This introduces a critical bias in performance evaluations, as methods that identify more pairs will have a higher chance of random overlap with known biological databases or results from other replicate datasets. Therefore, a fair comparison necessitates an adjusted framework that normalizes for these differences to account for this statistical artifact.

In addition to the statistical artifact of random overlaps, a fair evaluation is further complicated by a previously overlooked issue, the inherent expression bias within the biological databases that commonly used for validation. We observe that the known biological networks are biased toward genes and gene pairs with certain expression levels. To illustrate this, we calculated the average expression levels of the genes based on the lung dataset [24]. We contrast the expression levels of genes within biological networks versus the other genes and find that genes in biological networks are biased toward a higher expression level (Figure 8A and 8D). We further consider pair-level bias. We established the background expression level for gene pairs by calculating the geometric mean of the expression levels for every possible combination, using the genes present in the biological network. As shown in Figures 8B and 8E, the genes in the biological network, which are already biased in expression levels, exhibit an additional tendency toward higher expression at the pair level. Although there is a general tendency for genes and gene pairs in known biological networks to have higher expression levels, the degree of such bias varies across datasets. For example, while there is an expression bias across a broad range of genes for the lung dataset (Figure S11), the bias is negligible for the top 1,000 or 5,000 genes in the ROSMAP dataset (Figures S12 and S13). This dataset-specific nature of the bias means that a one-size-fits-all correction is inadequate and necessitates a more flexible approach. To address the biases from both random overlaps and expression levels, we developed a flexible adjustment strategy based on a stratified hypergeometric model. This model allows us to estimate the expected number of random overlaps and its standard deviation after accounting for expression-level bias, ultimately providing a standardized Z-score for evaluation. We first categorize gene pairs into k expression bins based on their mean expression levels. Within any given bin i, the number of overlaps that would occur by chance can be modeled precisely. Let *n*_*i*_ be the total number of gene pairs in bin i, *b*_*i*_ be the number of those pairs that are in the known biological network, and *c*_*i*_ be the number of identified correlated gene pairs. Then, the number of random overlap with the biological network in bin i follows a hypergeometric distribution, *Hypergeometric*(*n*_*i*_, *b*_*i*_, *c*_*i*_). The total expected random overlap and its variance can be calculated by summing the expectations and variances from all k bins as the sampling process is independent across bins. From this, we derive a Z-score which quantifies the significance of the observed overlap, corrected for both the total number of identified pairs and expression bias.

**Fig. 8.**
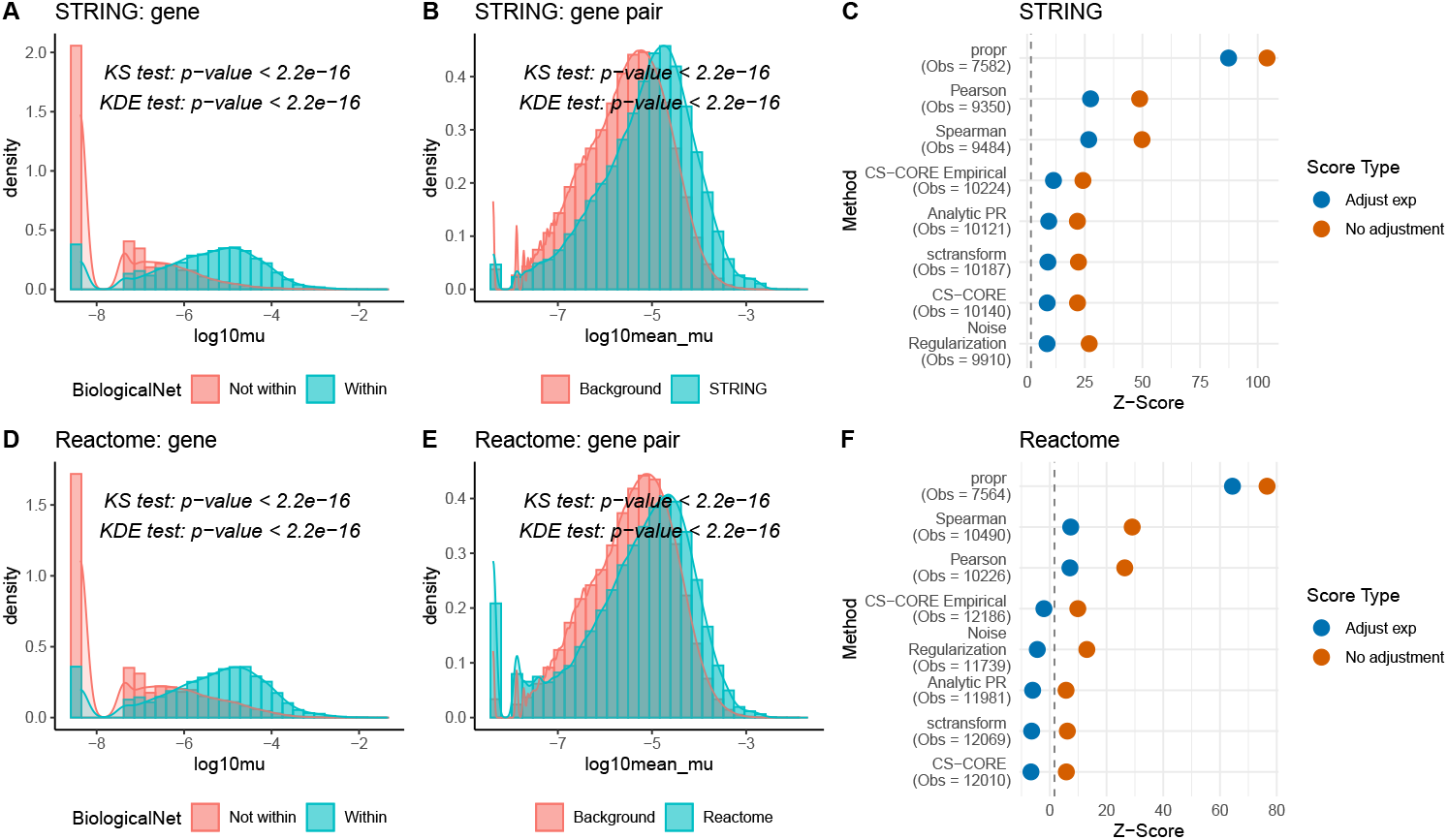
Expression bias in biological databases influences method performance rankings. We evaluated the expression bias in STRING (A-C) and Reactome (D-F). All analyses are based on monocytes from the lung dataset. **A and D**. Density plots comparing the mean expression of genes present in each network versus those not in the network. KS test and KDE test were conducted to assess the similarity between the two distributions, which demonstrate a significant gene-level bias. **B and E**. Density plots comparing the mean expression of gene pairs present in each network versus a background set, demonstrating a significant pair-level bias. **C and F**. Dot plots showing the Z-scores of various co-expression estimation methods with (Adjust exp) and without (No adjustment) the stratified expression adjustment. The number of observed overlaps with the biological database is shown in parentheses (Obs) for each method. The dashed line represents the significance threshold (one-sided p *<*0.05), which corresponds to a Z-score of 1.645.

Our adjustment strategy is designed to correct for two distinct sources of bias. Therefore, we demonstrate its importance in a two-step comparison. First, we compared the observed overlap counts of different methods with their calculated non-stratified Z-scores, which were generated without adjusting for expression level bias. As shown in Figures 8C and 8F, a naive comparison based on observed overlap counts can be misleading. For example, while CS-CORE with empirical p-values identifies the highest number of overlaps with known biological networks, its Z-score is not among the top-ranked. This occurs because a larger set of identified gene pairs inherently increases the chance of random overlaps, confirming that an adjustment for network size is crucial for a fair comparison.

We next demonstrate the necessity of our stratified approach for correcting expression-level bias by comparing its results to those from a non-stratified Z-score. When there is no apparent expression bias in a dataset, both the stratified and non-stratified approaches produce consistent method significance and rankings (Figures S14A and S14B). However, in the presence of strong expression bias, the stratification step becomes critical for an accurate evaluation. Applying our expression level adjustment has a significant impact on the evaluation outcomes. As shown in Figure 8F, the Z-scores for some methods become statistically insignificant, and the overall ranking of the methods changes. This demonstrates that the stratified approach is essential for disentangling a method’s true performance from the confounding effects of expression bias in validation databases.

The flexibility of our stratified framework allows it to be applied to other evaluation metrics, such as reproducibility. To demonstrate this, we compared the observed number of reproducible pairs between the ROSMAP and PNAS datasets with the Z-scores. The results again show that the raw counts can be misleading. While the Spearman method identifies a high number of reproducible pairs, its Z-score is comparable to that of other methods (Figure S14C). Furthermore, some co-expression estimation methods are inherently biased towards finding correlations among highly-expressed genes (Figure S15). Our stratified adjustment corrects for this by evaluating performance within different expression bins, providing a much fairer test. As seen with the Spearman which has a more obvious expression bias (Figure S15), its Z-score drops more after our adjustment is applied (Figure S14C).

## 3 Discussion

In this study, we first established that, for cell-type-specific analysis of scRNA-seq data, selecting correlated gene pairs based on p-values is a more robust and biologically meaningful approach than relying on correlation strength. This is because p-values, unlike correlation strengths, provide a statistical measure of confidence that accounts for both the effect size and the variability of the data. Our initial results confirm that a p-value-based selection consistently yields a higher proportion of true positives for CS-CORE. However, the practical application of this is limited, as many co-expression estimation methods either do not provide p-values or provide ones that are not well-controlled. To overcome this, we proposed scSimu, an accurate and efficient simulation method that can generate reliable empirical p-values for any co-expression estimation methods. This enabled us to demonstrate the superiority of the p-value-based approach across all estimation methods, and provide a universal and accessible framework to move beyond simple correlation strength and identify more robust correlated gene pairs.

Building on this p-value-based foundation, we next addressed the challenges in the fair method comparison. A simple comparison of overlaps is often misleading, as a method that identifies a larger number of gene pairs will have a higher chance of random overlaps. Furthermore, we illustrated the potential expression bias in the validation databases. To correct for both of these confounding factors, we proposed a stratified hyper-geometric model to ensure that the methods are evaluated by their true ability to detect co-expressions, rather than artifacts.

However, some limitations should be noted. First, the validation of scSimu was primarily performed on the droplet-based scRNA-seq data. Although we demonstrated its applicability to plate-based data like Smart-seq (Figure S17), a more extensive benchmark across a wider range of simulation methods, protocols and datasets would further evaluate its generalizability. Second, our empirical p-value approach, while robust, is computationally expensive. This may make it less practical for co-expression methods that already have extensive running times, for example, baredSC [25] and locCSN [26]. Third, our framework is not compatible with methods that rely on quantile normalization, such as SpQN[7], as this transformation alters the data distribution in a way that makes the simulated null data incomparable to the observed data. Finally, while our stratified model mitigates the impact of expression bias in validation databases, we still recommend considering diverse evaluation metrics as it is still unknown whether this bias is an artifact or represents real biology.

Ultimately, our study provides a rigorous workflow for co-expression analysis, moving from the initial principle of p-value superiority to a comprehensive framework for robust gene pair selection and unbiased method benchmarking. By providing both a practical guide and a new standard for evaluation, we hope to improve the reliability of co-expression analysis in the single-cell genomics field.

## 4 Methods

### 4.1 Datasets and preprocessing

Our study utilized three droplet-based datasets, including two single-nucleus RNA-sequencing (snRNA-seq) datasets of the human prefrontal cortex from the ROSMAP cohort [27] and a study by Lau et al. [28] (PNAS), and one scRNA-seq dataset from lung tissue [24]. To ensure consistency, we restricted our analysis to control subjects from each study. Detailed preprocessing steps are provided in Supplementary Note S1.

#### Biological networks

We obtained the human protein-protein interaction network from STRING v12 [11], filtering for interactions with a combined score greater than 500. The other biological network we considered was a pathway database, Reactome [12]. By combining each pair within the same pathway, we constructed an interaction network [1].

#### GO terms

The genes’ GO terms were obtained from the R package biomaRt version 2.58.0.

### 4.2 scSimu

For cell i and gene j, scSimu simulates the observed count, *x*_*ij*_, from the following distributions,

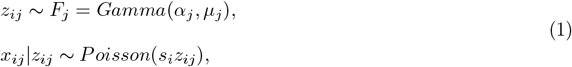

where *z*_*ij*_ is the corresponding relative underlying expression, *α*_*j*_ is the dispersion parameter, *µ*_*j*_ is the mean expression, and *s*_*i*_ is the sequencing depth. We model the dependence structure between genes at the relative underlying expression level using a copula, which can introduce the dependence among genes while preserving the marginal distributions. Based on Sklar’s theorem, the joint cumulative distribution function (CDF) of the relative underlying expression for p genes can be written as *F* (*z*_*i*1_, …, *z*_*ip*_) = *C*(*F*_1_(*z*_*i*1_), …, *F*_*p*_(*z*_*ip*_)), where *C*(·) is a copula. By employing a Gaussian copula with a target gene correlation matrix R, this relationship is defined specifically as *F* (*z*_*i*1_, …, *z*_*ip*_) = Φ_*R*_(Φ^−1^(*F*_1_(*z*_*i*1_)), …, Φ^−1^(*F*_*p*_(*z*_*ip*_))), where Φ_*R*_ is the CDF of multivariate normal distribution with mean vector zero and covariance matrix equal to the correlation matrix R, and Φ^−1^ is the inverse CDF of standard normal distribution. Therefore, we can use an inverse procedure to generate a multivariate distribution with the specified correlation structure and marginal distributions.

Compared with the simulation method used in our previous publication [9], here we updated the Gaussian copula process to allow for the case that the correlation matrix is not positive semi-definite given that some gene co-expression estimation methods sacrifice positive semi-definite to accommodate the noise and sparsity in scRNA-seq (e.g., CS-CORE, SpQN). Additionally, after applying a threshold based on significance or correlation strength, a positive semi-definite correlation matrix may no longer remain positive semi-definite. Our updated Gaussian copula process addresses this by employing eigendecomposition to replace any negative eigenvalues with zero. This procedure provides a simulation with a correlation structure that is more similar to the true one compared to our previous method of iteratively adding small values to the diagonal (Figure S16). In addition, unlike methods such as SymSim [29], ZINB-WaVE, powsimR [30], and muscat [31], scSimu preserves all original genes and cells in the output which is critical for accurate downstream comparisons.

The detailed simulation procedures and the zero-inflated model are described in Supplementary Note S2.

### 4.3 Benchmark simulation methods

To evaluate the performance of different simulation methods, we compared the simulated data with real data from eight data summaries:

- **Mean(logCPM)**: mean expression of genes estimated via log-normalized data.
- **Var(logCPM)**: variance of genes estimated using log-normalized data.
- **Coefficient of variation**: 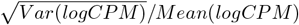.
- **Zero fraction (cell)**: fraction of cells with zero counts.
- **Zero fraction (gene)**: fraction of genes with zero counts.
- **Library size**: total gene counts per cell.
- **Cell correlation**: Spearman correlation between cells based on log-normalized data. The number of cells used to estimate cell correlation was capped by 500.
- **Gene correlation**: correlations estimated by CS-CORE for the top 1,000 highly expressed genes determined by Mean(logCPM).

To quantify the similarity between simulated and real data distributions, we employed both Kolmogorov-Smirnov (KS) and Kernel Density Estimation (KDE) tests. These tests offer complementary insights, with the KS test measuring the largest discrepancy between distributions and the KDE test assessing overall divergence. To ensure comparability across different datasets and metrics, KS and KDE statistics were min-max normalized separately by dataset and metric, then averaged to combine them. The overall performance of each simulation method was evaluated based on the mean score across eight data metrics. Comprehensive details and the benchmark for Smart-Seq data simulation are presented in Supplementary Note S3.

### 4.4 Empirical p-value estimation

The empirical p-values were estimated in three steps. (1) Generate k independent datasets corresponding to the real data using scSimu and estimate the correlations. For each gene pair, gather k estimates from these independent datasets alongside one from the real data. (2) Estimate the mean *µ*_0_ and variance 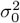of these k estimates for each gene pair. (3) The empirical p-value is calculated using a normal distribution with the estimated mean and variance, where the p-value = *P* (|*X*| *>* |*Estimate*_*obv*_|) and 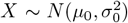. See Supplementary Note S4 for the analysis of type I error control.

### 4.5 Evaluation of estimated gene co-expression networks

To compare the performance of p-value-based and correlation-strength-based approaches, we generated synthetic datasets with known ground truth based on oligodendrocytes from ROSMAP and PNAS control subjects. We created a sparse ground truth by setting weak correlations to zero in the real data and simulating new counts using scSimu. We then estimated co-expression networks for the top 1,000 highly expressed shared genes using seven different methods. More details can be found in Supplementary Note S5.

#### Reproducibility

We used the simulated PNAS data as independent validation data. The reproducibility was evaluated based on the number of reproducible pairs which are gene pairs identified as correlated in both datasets.

#### Precision

Precision is the ratio of true positives to inferred positives. For reproducibility, the precision values in the main text were based on the ROSMAP data, while those from the PNAS data were displayed in Figures S1 and S6.

#### Proportion of misidentification

Let *A* and *B* be the sets of identified correlated gene pairs from two datasets, with *A*_*T*_ and *B*_*T*_ representing their respective true positives. The proportion of misidentified reproducible pairs is calculated as (|*A* ∩ *B*| − |*A*_*T*_ ∩ *B*_*T*_ |)*/*|*A* ∩ *B*|. Similarly, for a biological network *C*, the proportion of misidentified overlaps is (|*A* ∩ *C*| − |*A*_*T*_ ∩ *C*|)*/*|*A* ∩ *C*|.

### 4.6 Expression bias in biological networks

To demonstrate the expression bias in the STRING and Reactome, we estimated the mean expression level for each gene using monocytes from the lung dataset and a Gamma-Poisson GLM. We first assessed the gene-level bias by comparing the expression distributions of genes in these biological networks versus those that are not. We then focused on genes within the networks to evaluate pair-level bias. We further investigated the bias by repeating the pair-level analysis on subsets of genes, including the top 100, 1,000, and 5,000 most highly expressed or highly variable genes. For each of these analyses, the biological network was correspondingly filtered to include only interactions among the selected top k genes. The background was generated based on this sub-network. To confirm that these findings were not dataset-specific, we also studied the observed bias using mean expression levels from oligodendrocytes in the ROSMAP.

### 4.7 A stratified hypergeometric model for bias adjustment

First, all gene pairs are categorized into k bins based on their mean expression level. The number of random overlaps is then modeled by a separate hypergeometric distribution within each bin i, *Hypergeometric*(*n*_*i*_, *b*_*i*_, *c*_*i*_). The total expected number of random overlaps and its variance are calculated by summing the expectations and variances from all k bins, 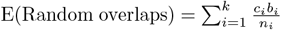 and 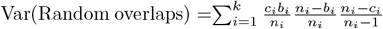. This yields a stratified Z-score corrected for both network size and expression bias. We validated the stratified framework using the lung dataset by assessing the overlap of identified networks with the STRING and Reactome databases. Details regarding the baseline non-stratified model and the application to real data are provided in Supplementary Note S6.

## Declarations

### Availability of data and materials

The ROSMAP dataset (syn52293417) has controlled access under the regulation of human privacy. The PNAS dataset used in our analysis is publicly available (GSE157827). The lung data we used to evaluate the expression bias in biological networks can be accessed through GSE136831. The Smart-Seq data can be accessed through the Single Cell Portal with accession numbers SCP424. The relevant code of this study can be found in https://github.com/xs222/coexp validation [32]. The R package scSimu is available at https://github.com/xs222/scSimu [33].

### Authors’ contributions

XS conducted the study. YL participated the discussion of findings and provided feedback on the manuscript draft. HZ supervised the project. All authors read and approved the final manuscript.

## Acknowledgements

We thank Dr. Jia Zhao for her insightful feedback and helpful discussions. We also would like to thank the Religious Orders Study and Memory and Aging Project (ROSMAP) Study, Dr. Shun-Fat Lau, Dr. Nancy Y. Ip, Taylor S. Adams, Dr. Naftali Kaminski, Dr. Ding Jiarui and Dr. Joshua Z. Levin for sharing the datasets.

## Supplementary information

### Supplementary Note S1: Datasets and preprocessing

As a preprocessing step, we selected only the control subjects from each study, which included seven subjects with a Braak Stage of 0 (ROSMAP), nine healthy controls (PNAS), and 28 control subjects (lung). We used the gene expression count data for all datasets and the cell-type labels for the ROSMAP and lung datasets directly from their original studies. Given that the cell-type annotations for PNAS data are not available from the original paper, we adopted the annotations provided by Su et al. [9], which were generated using the procedure outlined in the original study. The gene number and cell number for all cell types used in our analyses are detailed in Table S3.

### Supplementary Note S2: scSimu model details

#### Choices of model parameters

The input for the simulation framework can be a count matrix or userdefined parameters. For count matrix input, we use the observed sequencing depths (*s*_*i*_), estimate the marginal parameters for genes (*α*_*j*_ and *µ*_*j*_) based on the Gamma-Poisson generalized linear model (GLM) [34], and calculate the correlation matrix for genes that are highly expressed, highly variable, or of particular interest by CS-CORE. Alternatively, users may opt to directly provide marginal parameters, sequencing depths, and correlation matrix based on their preferences.

#### Detailed procedures

For simulation, we mimic the sequencing procedure commonly adopted. We first generate a gene expression matrix under the Gamma distribution. If the correlation matrix is not an identity matrix, i.e., genes are correlated, the correlation structure inferred from real data or specified by the user will be imposed through the Gaussian copula. Then, the gene count matrix is simulated from the underlying gene expression matrix using the Poisson distribution with the sequencing depths from the input real data or the user.

#### Zero-inflated simulation

Additionally, we extended the framework to simulate zero-inflated data. This extends the applicability of our method to protocols beyond droplet-based technologies. We assume that zero-inflation is a technical artifact distinct from biological zeros, as discussed by Jiang et al. [22]. The simulation process begins by classifying each gene as either zero-inflated or not using a likelihood ratio test. Next, we use a Gamma-Poisson GLM for non-zero-inflated genes and a zero-inflated model for zero-inflated genes, which also provides the zero-inflation parameter (p), to estimate the marginal parameters. We then generate a count matrix for all genes from the negative binomial distribution described above. Finally, we introduce additional zeros into the zero-inflated genes by randomly sampling cells to be zero, with the sampling probability equal to the estimated zero-inflation parameter p. We validated this zero-inflation framework on Smart-Seq data, where scSimu achieved significantly better performance than scDesign3, confirming its robustness across different scRNA-seq protocols (Figure S17). More details can be found in Supplementary Note S3.

### Supplementary Note S3: Benchmark simulation details

#### Benchmark datasets

We used two datasets in our comparisons, the prefrontal cortex of seven subjects with a Braak Stage of 0 from Mathys et al. [27] (ROSMAP) and the prefrontal cortex of nine healthy controls from Lau et al. [28] (PNAS). From each of these two cohorts, we analyzed five cell types resulting in ten distinct datasets for our evaluation. To save computation time, cell types with more than 10,000 cells were randomly downsampled to 5,000. The details for each dataset are provided in Table S3.

#### Implementation of simulation methods

We considered 13 simulation methods in addition to scSimu. For the independent task, we considered nine other methods (Table S1). ZINB-WaVE, SymSim, POWSC, and Hierarchicell were excluded because they are not able to generate independent genes. We did not compare with the permutation method proposed by Specht and Li [35] as it permutes normalized data and cannot be applied to co-expression estimation methods that require a count matrix as input. For correlated task, we compared scSimu with 11 other simulation methods (Table S1). Although ESCO can incorporate correlation structures, it is unsuitable for targeted correlation analysis because it selects the gene pairs to be correlated at random. We also did not consider SPsimSeq [36] for both tasks because of the extensive computational time required by this method. For example, for a dataset with 33,538 genes and 4,724 cells, SPsimSeq took 2.5 hours to simulate independent genes, and 30 hours to simulate correlated genes. The descriptions of these simulation methods are listed below. The input for all benchmarked methods is a SingleCellExperiment object containing the real count data. For any parameters not explicitly mentioned in this section, we used the default settings for that method.

- **ESCO**: Independent genes were simulated using functions from R package ESCO version 0.99.12 by specifying `withcorr=F` and `type = “single”`.
- **hierarchicell**: Conducted using R package hierarchical version 1.0.0.
- **muscat**: Conducted through R package muscat version 1.14.0.
- **Permutation**: Followed procedures in Su et al. [9]. Specifically, this method first converts the raw count matrix into expression ratios by dividing the count of each cell by its sequencing depth. To break the correlation structure, the expression ratios for each gene are then randomly shuffled across all cells. Finally, we generate a new count matrix by sampling from a Poisson distribution, using the product of the shuffled gene ratios and the observed sequencing depths as the mean.
- **POWSC**: Simulated using R package POWSC version 0.99.12.
- **powsimR**: It simulated data containing both case and control cells based R package powsimR version 1.2.5 as originally designed for differential expression analysis.
- **scDesign2**: Simulation was conducted using R package scDesign2 version 0.1.0. Set `marginal=“nb”` to reduce computational time, while default settings were retained for all other parameters. Independent genes were simulated by specifying `sim_method = “ind”`.
- **scDesign3**: We set the mean formula to be `mu_formula = “`offset(log(library size))”.
- Correlated genes were simulated by setting `corr_formula = “1”` and independent genes were simulated by setting `corr_formula = “ind”` utilizing R package scDesign3 version 1.0.1.
- **SCRIP**: Simulation were conducted by using R package SCRIP version 1.0.0.
- **SPARSim**: Simulated using R package sparsim version 0.9.5.
- **Splat**: Conducted by using R package splatter version 1.26.0.
- **SymSim**: Performed using the R package SymSim version 0.0.0.9000.
- **ZINB-WaVE**: We utilized the function related to ZINB-WAVE in the splatter package to simulate data.

#### Evaluation metrics

The KS test statistics were calculated via the function `ks.test` in the R package stats version 4.3.0. The KDE test statistics were calculated by`kde.test` function in the R package ks version 1.14.1. For metrics exceeding 10,000 points (e.g., gene correlation), we only used 10,000 of them to calculate the KDE test statistics to reduce the computation time. All computational time benchmarks were assessed via the Intel Xeon Platinum 8358 and Intel Xeon Gold 6240 processors.

#### Benchmark methods for simulating Smart-Seq data

To evaluate the performance of scSimu on Smart-Seq data, we compared it with scDesign3 using a dataset from Ding et al. [37]. We preprocessed this data by subsetting to Cytotoxic T cells from the PBMC1 experiment and then removing genes that were expressed in fewer than 10 cells or had a total count below 10. For the scDesign3 simulation, we specified a zero-inflated negative binomial marginal distribution by setting `family use = “zinb”`.

### Supplementary Note S4: Type I errors

#### Type I errors

To demonstrate that all gene co-expression estimation methods can achieve well-controlled type I errors using such empirical p-values, we first used scSimu to mimic oligodendrocytes in the PNAS data. We calculated the empirical p-values of estimated correlations for the top 1,000 highly expressed genes based on the procedure above with *k* = 500. We then compared the empirical p-values of independent gene pairs with Unif(0,1) in Figure 3.

### Supplementary Note S5: Evaluation of estimated gene co-expression networks details

#### Real data simulation

To compare the p-value-based and correlation-strength-based approaches, we used simulated data with known ground truth. This simulation was based on oligodendrocytes from controls in the ROSMAP and PNAS datasets. We first normalized the real data using sctransform and calculated a Pearson correlation matrix for the 1,000 most highly expressed shared genes. To create a sparse and well-defined ground truth, we set weak correlations to zero (abs(cor) *<* 0.015 for ROSMAP, abs(cor) *<* 0.017 for PNAS). Then, we used scSimu to simulate new count data based on these modified correlation matrices. This gives us two synthetic datasets, one mimicking the ROSMAP data and the other mimicking the PNAS data. Each of these datasets has known gene correlation structures, providing a ground truth for our comparisons.

#### Estimation of gene co-expression networks

Using the simulated data, we estimated the co-expression networks for the top 1,000 genes with the highest expression levels shared between the ROSMAP and PNAS datasets using seven methods. The Analytic PR inferred the gene co-expression network using Pearson correlation on the residuals obtained through the procedures in Lause et al. [23]. CS-CORE was implemented through functions in R package CS-CORE version 0.0.0.9000. Noise Regularization was based on the scrips in https://github.com/RuoyuZhang/NoiseRegularization [38]. Pearson and Spearman estimated the gene coexpression networks by the Pearson and Spearman correlation on log-normalized data, respectively. The R package propr version 4.2.6 was used to generate the gene co-expression networks for propr. For sctransform, we used R package Seurat version 5.0.1 to derive the residuals and then applied Pearson correlation to them.

#### Approaches for selecting correlated gene pairs

We evaluated and compared the correlation-strength-based approach and p-value-based approach for selecting correlated gene pairs.

- **Correlation-strength-based approach**. We defined correlation strength as the absolute value of the estimated correlation. The correlated gene pairs are the top k gene pairs with the largest correlation strength. The k was varied from 1,000, 5,000, 10,000, 50,000, to 100,000. For reproducibility, we included the number of true correlated gene pairs for the ROSMAP simulated data (135,244) and PNAS simulated data (59,933) as our cutoffs. For the evaluation of overlaps with known biological networks, we included the number of true correlated gene pairs for the ROSMAP simulated data (135,244) [4].
- **P-value-based approach**. The p-values were adjusted according to the Benjamini-Hochberg (BH) procedure. The correlated gene pairs are those pairs with adjusted p-values less than the threshold. The thresholds for the p-values considered here are 0.001, 0.005, 0.01, 0.05, and 0.1.

### Supplementary Note S6: Hypergeometric model for bias adjustment

**Table S1.**
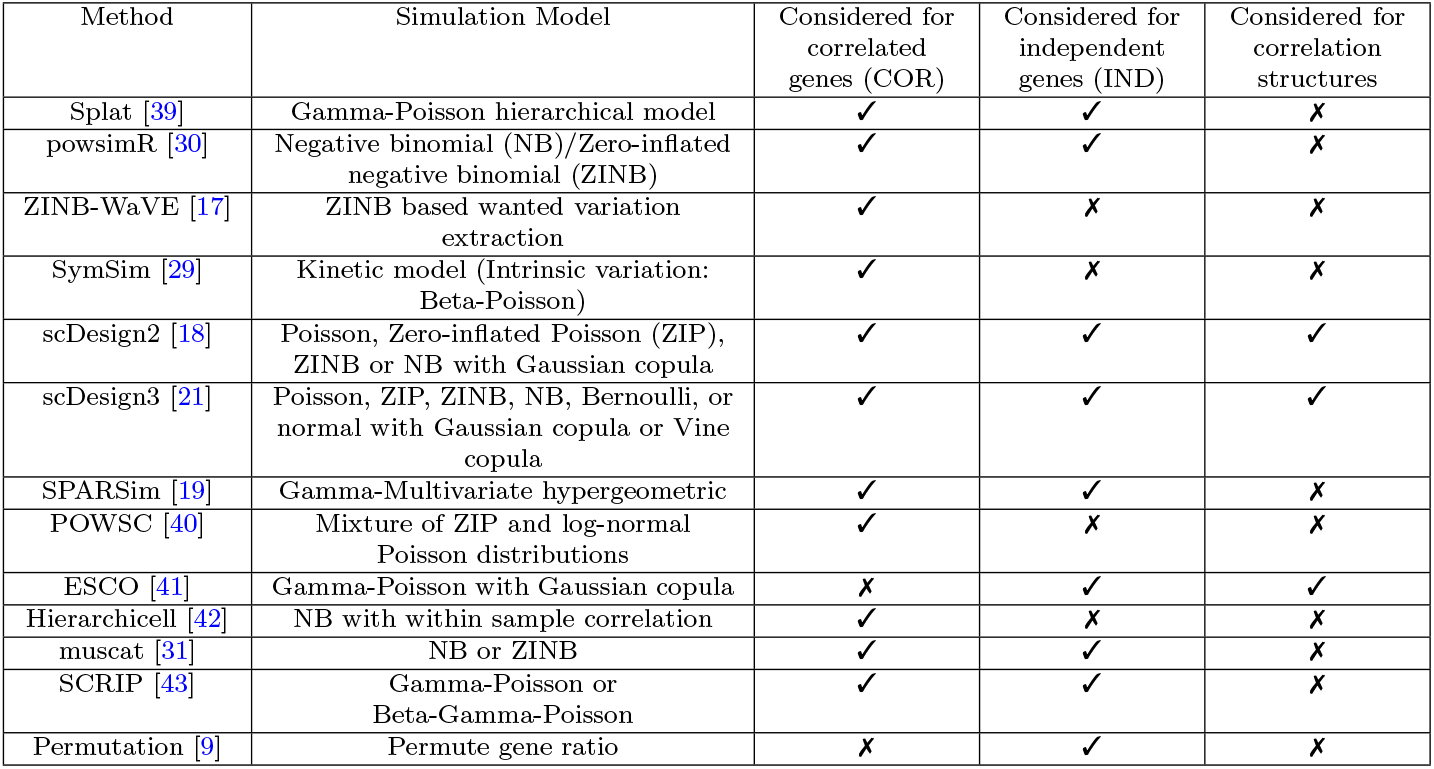
Simulation methods compared in our evaluation.

**Table S2.** Proportion of non-zeroes among the off-diagonal entries in the independent simulation.

| Method | Proportion of non-zeroes | Method | Proportion of non-zeroes |
| --- | --- | --- | --- |
| scSimu | 0 | Permutation | 0 |
| ESCO | 0 | muscat | 0.0287 |
| SCRIP | 0 | powsimR | 0.0007 |
| SPARSim | 0.0006 | scDesign2 | 0 |
| Splat | 0 | scDesign3 | 0 |

**Table S3.** Summary of real data used in the analysis.

| Dataset | Tissue | Protocol | Data type | # genes | Cell types used for benchmark (# cells) |
| --- | --- | --- | --- | --- | --- |
| ROSMAP [27] | Prefrontal cortex | 10x Genomics | snRNA-seq | 33538 | Astrocytes (2551),<br>Excitatory neurons (4724),<br>Immune cells (1198),<br>Oligodendrocytes (15518),<br>Oligodendrocyte precursor cells (1545) |
| PNAS [28] | Prefrontal cortex | 10x Genomics | snRNA-seq | 28412 | Astrocytes (7767),<br>Endothelial (688),<br>Excitatory neurons (24429),<br>Inhibitory neurons (5976),<br>Oligodendrocytes (17693) |
| Lung [24] | Lung | 10x Genomics | scRNA-seq | 45947 | Monocytes (5989) |

**Fig. S1.**
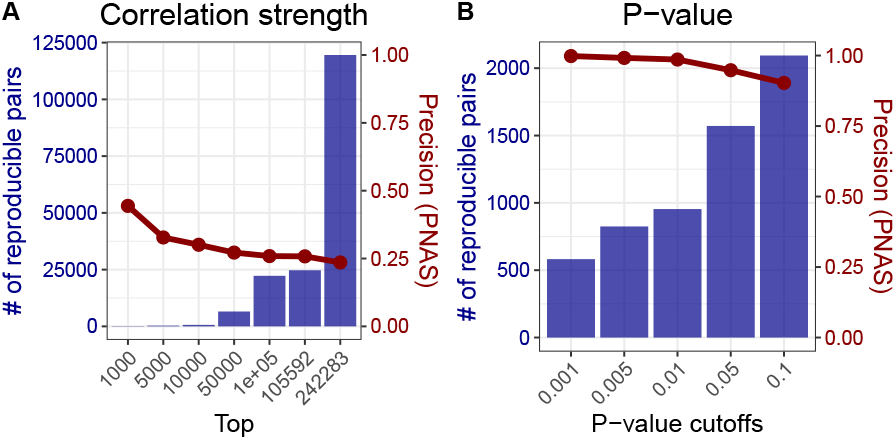
Precision in PNAS data versus the number of reproducible pairs for CS-CORE. Correlated gene pairs were identified using the correlation-strength-based approach (**A**), and p-value-based approach (**B**).

**Fig. S2.**
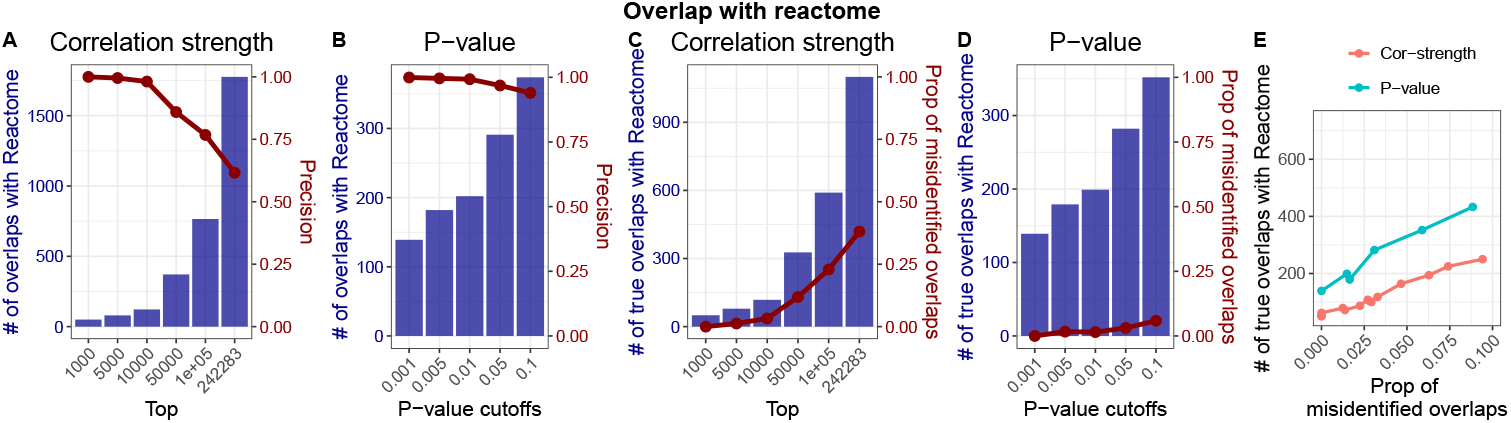
Comparison of correlation-strength-based and p-value-based approaches in terms of overlaps with Reactome using CS-CORE. **A-B**. Total number of overlaps with the Reactome and precision for the correlation-strength and p-value (B) approaches. **C-D**. Number of true overlaps with Reactome and the proportion of misidentified overlaps for the correlation-strength (C) and p-value (D) approaches. **E**. Direct comparison of the number of true overlaps versus the proportion of misidentified overlaps.

**Fig. S3.**
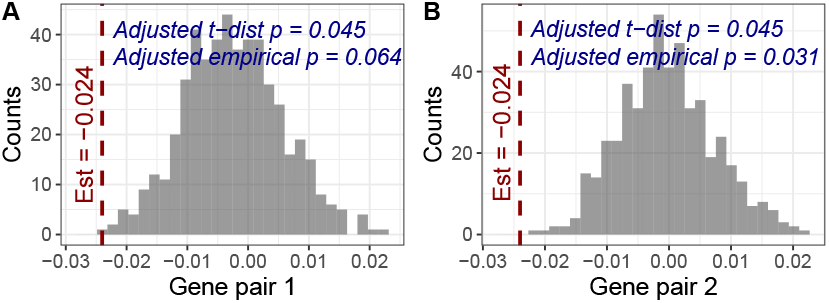
P-values from Student’s t-distribution may not reflect the underlying null distribution. Gene pairs were selected from the simulation used in Section 2.1 where gene pair 1 is independent and gene pair 2 is correlated. The figures display the histograms of their correlation estimates under the independent simulations, with the observed estimate indicated by a dashed dark red line. The adjusted two-sided t-distribution and empirical p-values are displayed in dark blue. Importantly, the adjusted t-distribution p-values are the same, failing to distinguish between the independent and correlated cases. In contrast, the adjusted empirical p-values correctly capture this difference, showing a non-significant result for the independent pair (adjusted p=0.064) and a significant result for the correlated pair (adjusted p=0.031).

**Fig. S4.**
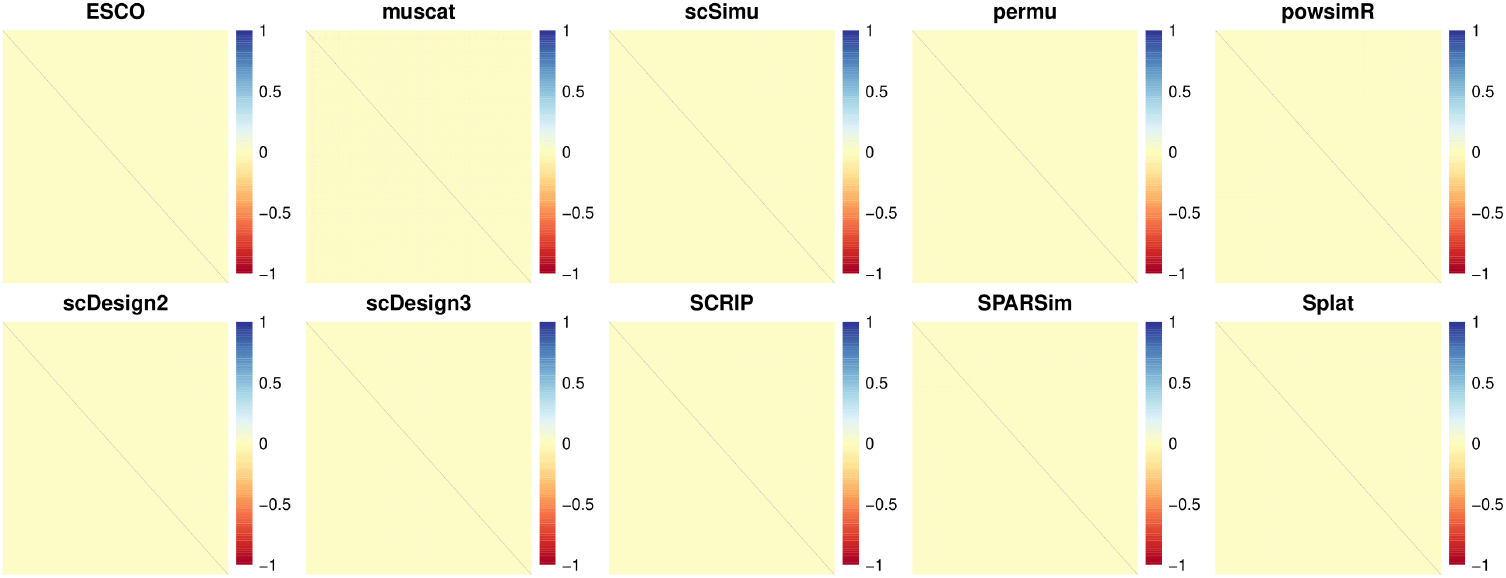
Independence of the simulated independent data of real data. Heatmap for correlation matrix of the top 1,000 highly expressed genes under the independent simulation.

**Fig. S5.**
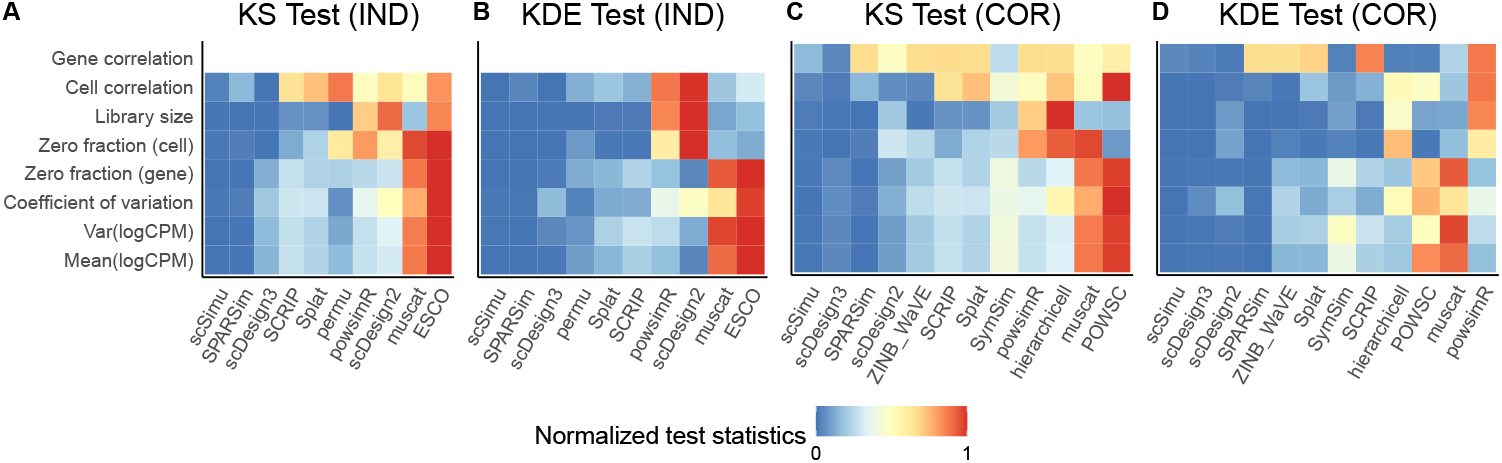
Benchmark of the simulation methods using KDE test and KS test separately. The x-axis is the simulation method and the y-axis is the evaluation metric. Color represents the normalized test statistics of the KS test or KDE test. The x-axis is ordered based on the mean of the normalized test statistics. **A and B**. Evaluate the simulation of the corresponding independent data of real data based on the KS test and KDE test respectively. **C and D**. Evaluate the simulation of real data based on the KS test and KDE test respectively.

**Fig. S6.**
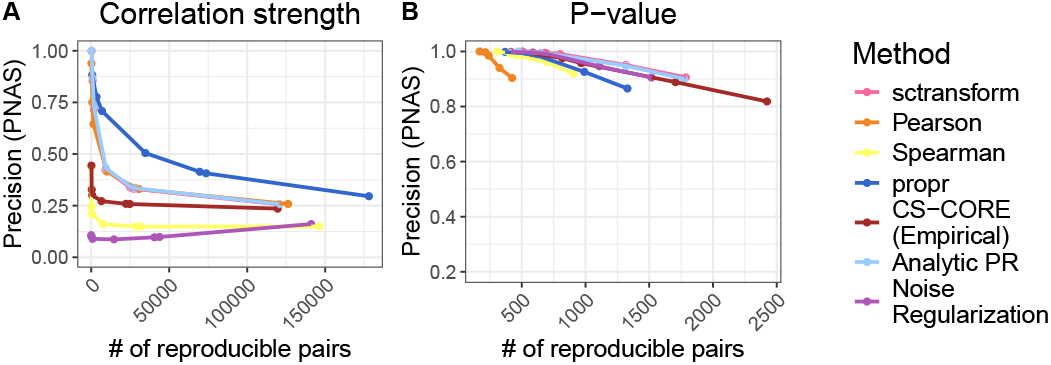
Precision in PNAS data versus the number of reproducible pairs. Correlated gene pairs were identified using the correlation-strength-based approach (**A**), and p-value-based approach (**B**).

**Fig. S7.**
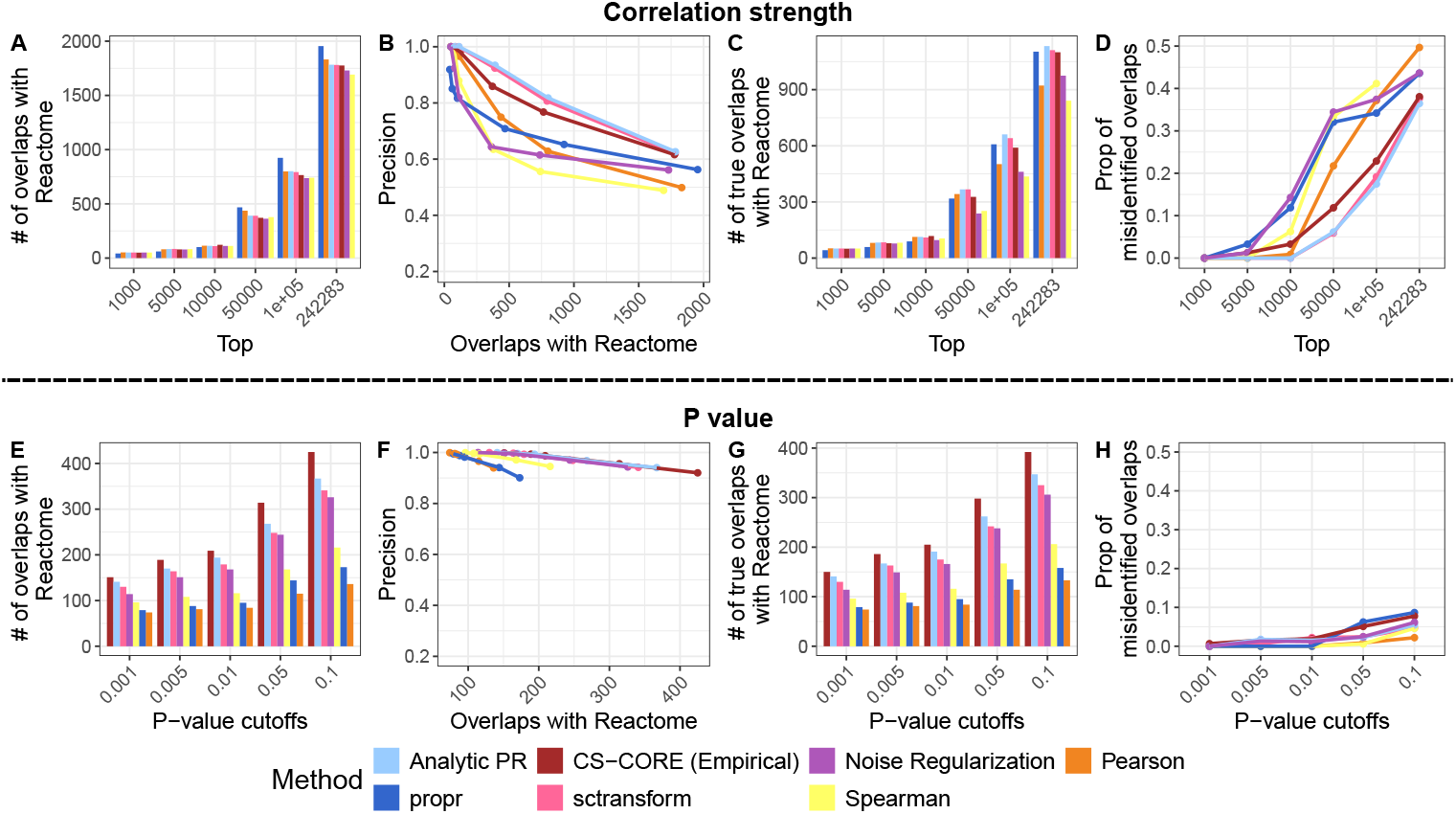
Performance in terms of overlaps with Reactome. **A. and E**. The number of overlaps with Reactome at different cutoffs. **B and F**. Relationship between precision and the number of overlaps with Reactome. **C and G**. The number of true overlaps with Reactome at different cutoffs. **D and H**. The proportion of misidentified overlaps when gene pairs were selected by their estimated correlations (D) or by the empirical p-values (H).

**Fig. S8.**
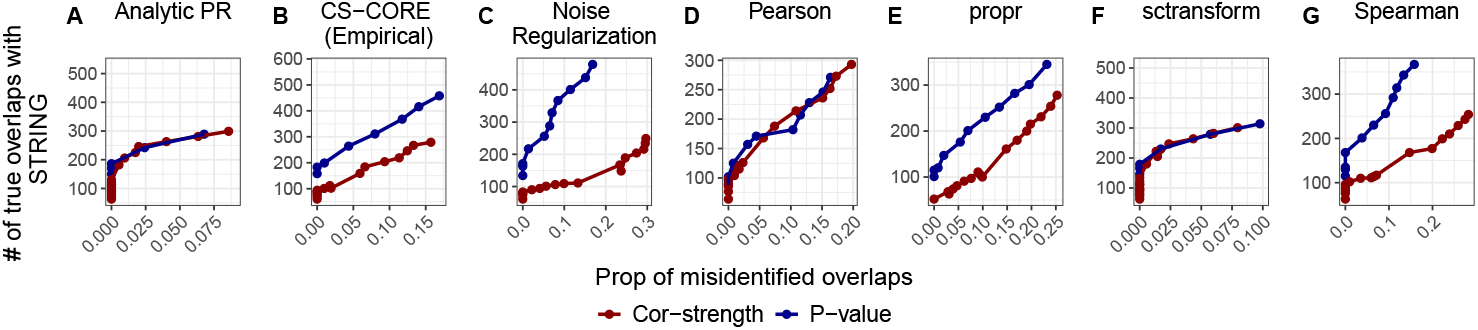
Comparison of correlated gene pairs selection approaches based on the overlaps with STRING. The performance of selecting gene pairs by correlation strength (dark red) and p-value (dark blue) was evaluated for seven different co-expression estimation methods. Each panel plots the number of true overlaps with STRING (y-axis) against the proportion of misidentified reproducible pairs (x-axis) at varying selection thresholds.

**Fig. S9.**
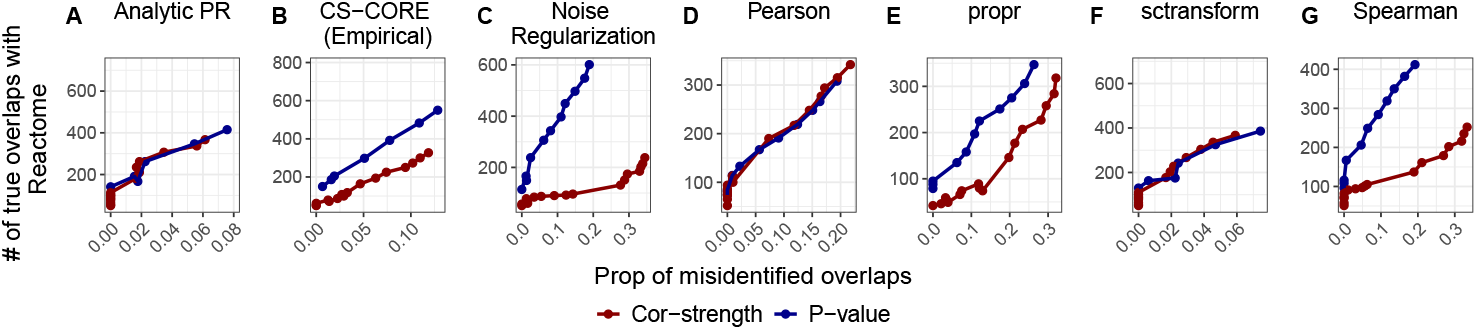
Comparison of correlated gene pairs selection approaches based on the overlaps with Reactome. The performance of selecting gene pairs by correlation strength (dark red) and p-value (dark blue) was evaluated for seven different co-expression estimation methods. Each panel plots the number of true overlaps with Reactome (y-axis) against the proportion of misidentified reproducible pairs (x-axis) at varying selection thresholds.

**Fig. S10.**
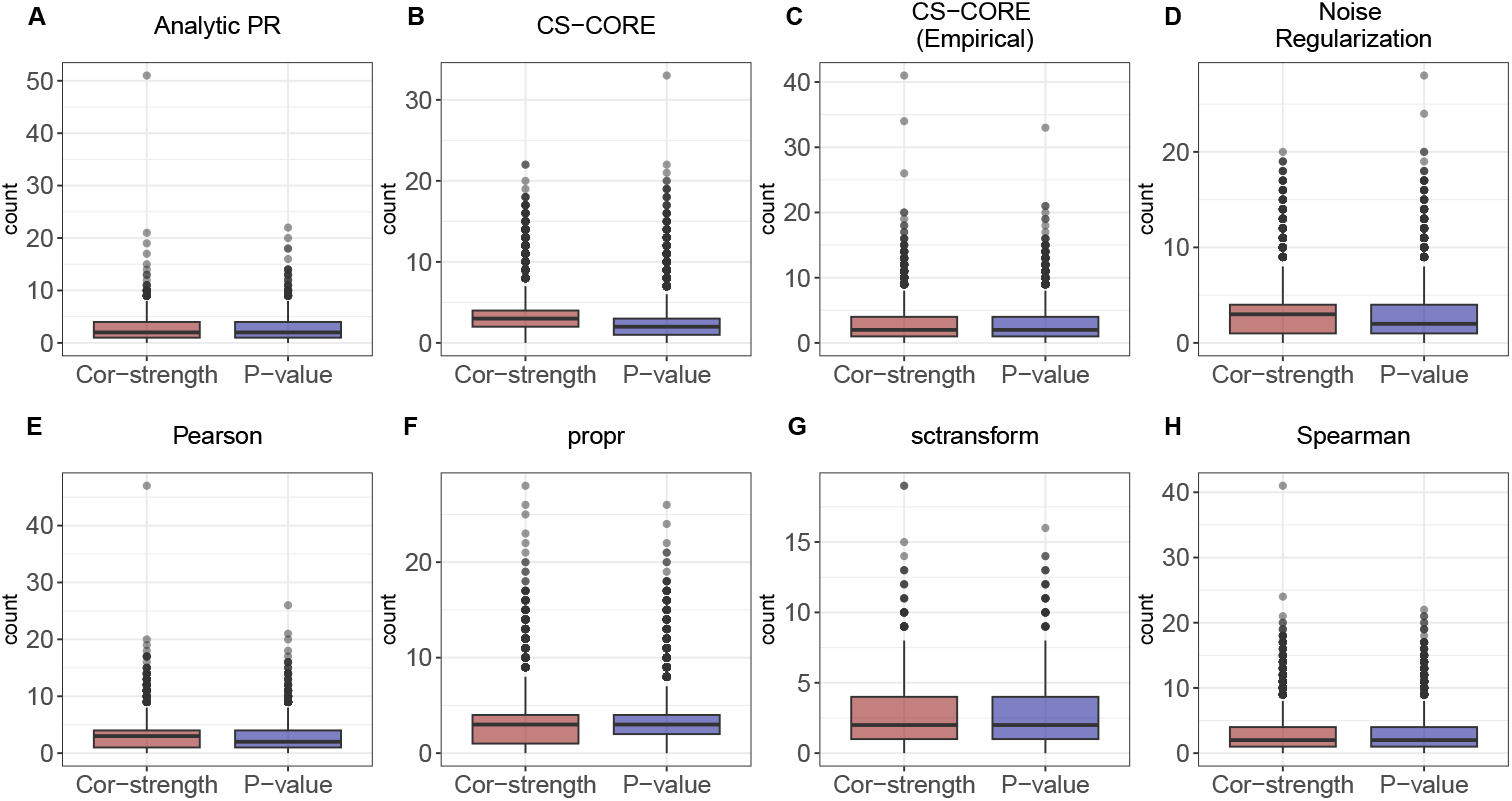
Comparison of shared GO terms in oligodendrocytes. For each estimation method, we identified two sets of gene pairs in oligodendrocytes, those found only by the p-value-based approach and those found only by the correlation-strength-based approach. To ensure a fair comparison, the threshold for the correlation-strength approach was set to yield the same number of gene pairs as the p-value approach at a 0.05 significance level (BH adjusted). The boxplots display the distribution of the number of shared GO terms for the gene pairs in each exclusive set.

**Fig. S11.**
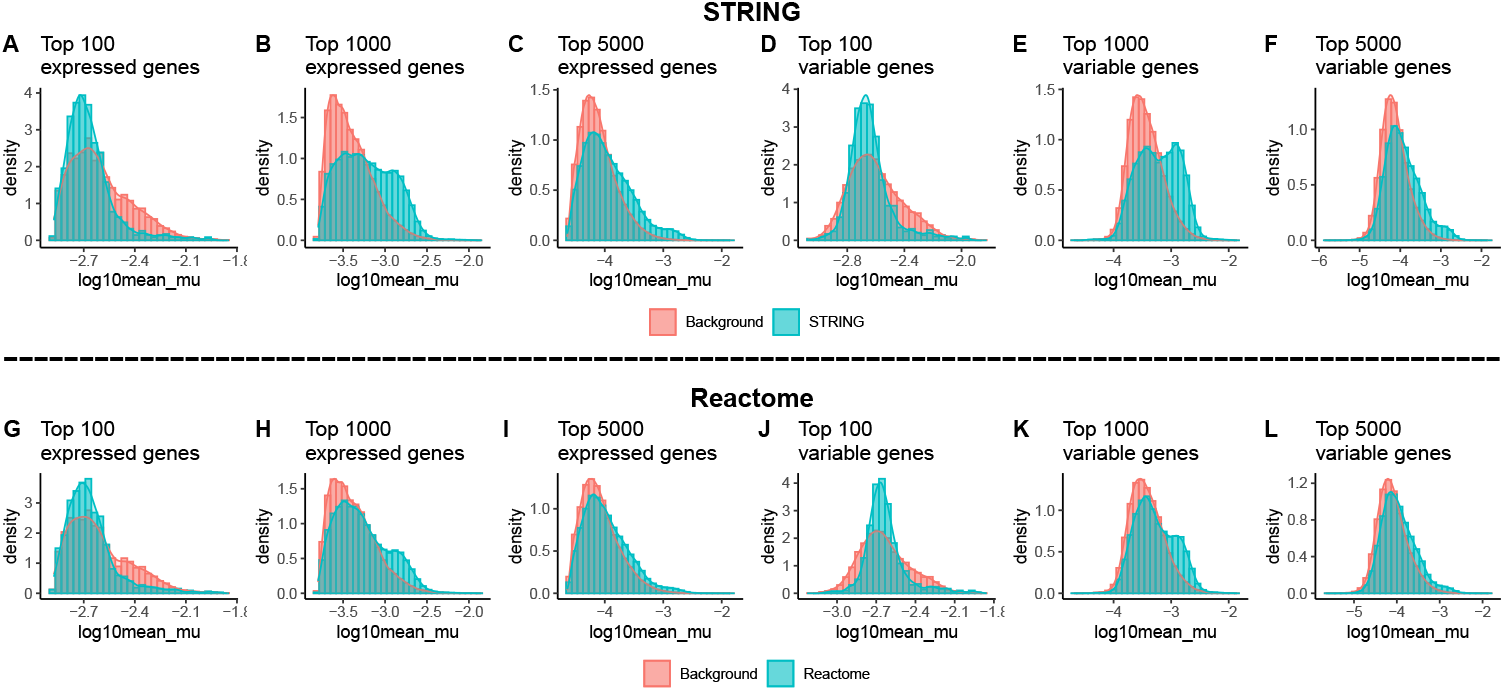
Expression bias in the STRING and Reactome databases for a subset of genes. We evaluated the expression bias in STRING (A-F) and Reactome (G-L) for a subset of genes based on monocytes from the lung dataset. Each panel shows a density plot for comparing the mean expression of gene pairs present in a subset of the biological databases versus the corresponding background. See Section 4.6 for full procedural details.

**Fig. S12.**
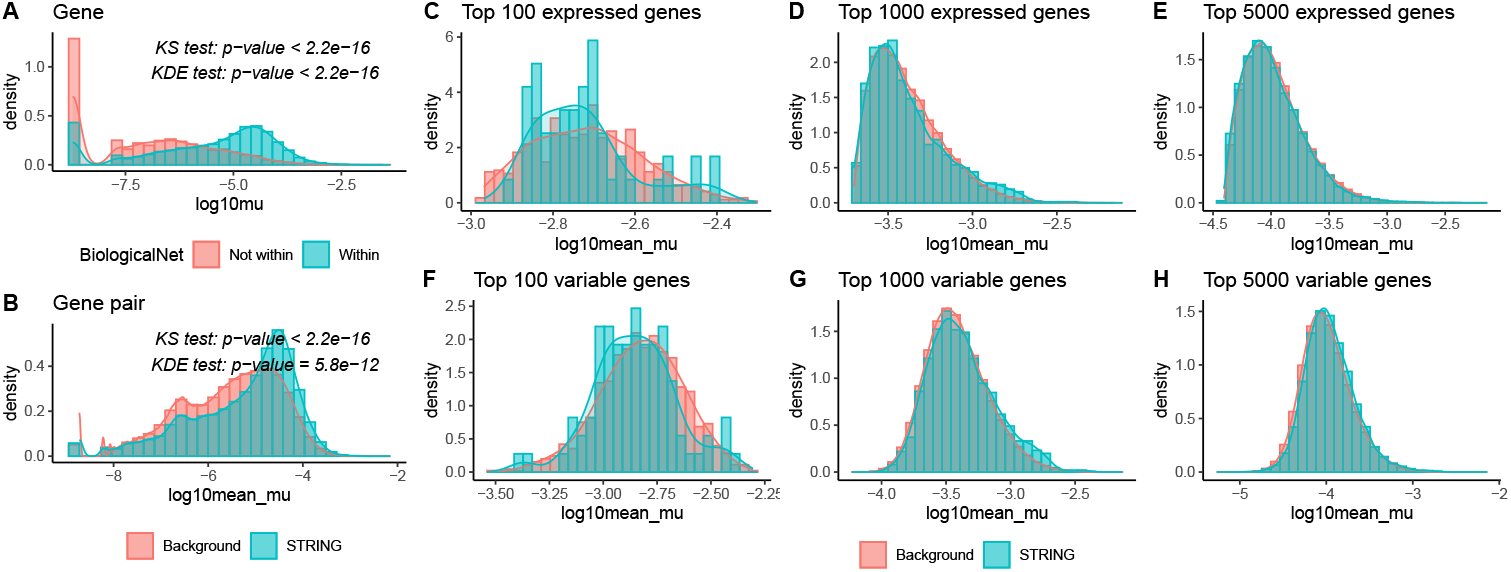
Expression bias in STRING where gene expression levels were derived from the ROSMAP oligodendrocytes. **A**. Density plot comparing the mean expression of genes present in STRING versus those not in the network. **B**. Density plot comparing the mean expression of gene pairs present in STRING versus a background set. **C-H**. Density plots comparing the mean expression of gene pairs present in a subset of STRING versus the corresponding background.

**Fig. S13.**
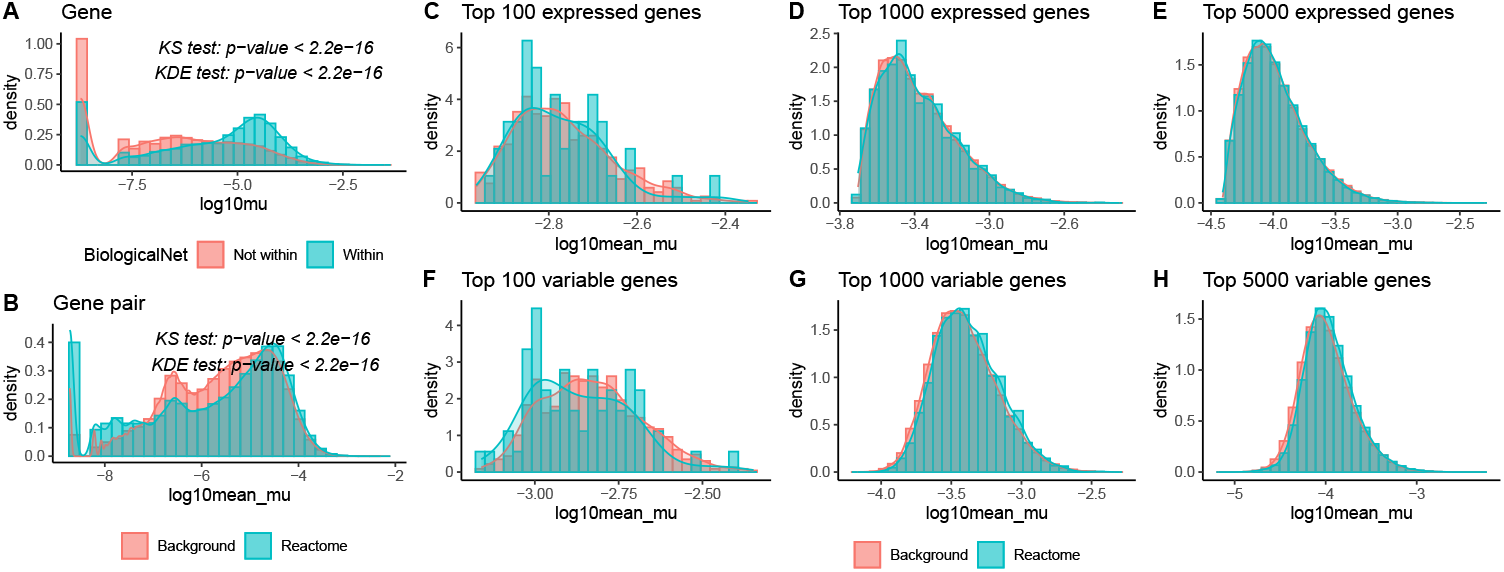
Expression bias in Reactome where gene expression levels were derived from the ROSMAP oligodendrocytes. **A**. Density plot comparing the mean expression of genes present in Reactome versus those not in the network. **B**. Density plot comparing the mean expression of gene pairs present in Reactome versus a background set. **C-H**. Density plots comparing the mean expression of gene pairs present in a subset of Reactome versus the corresponding background.

**Fig. S14.**
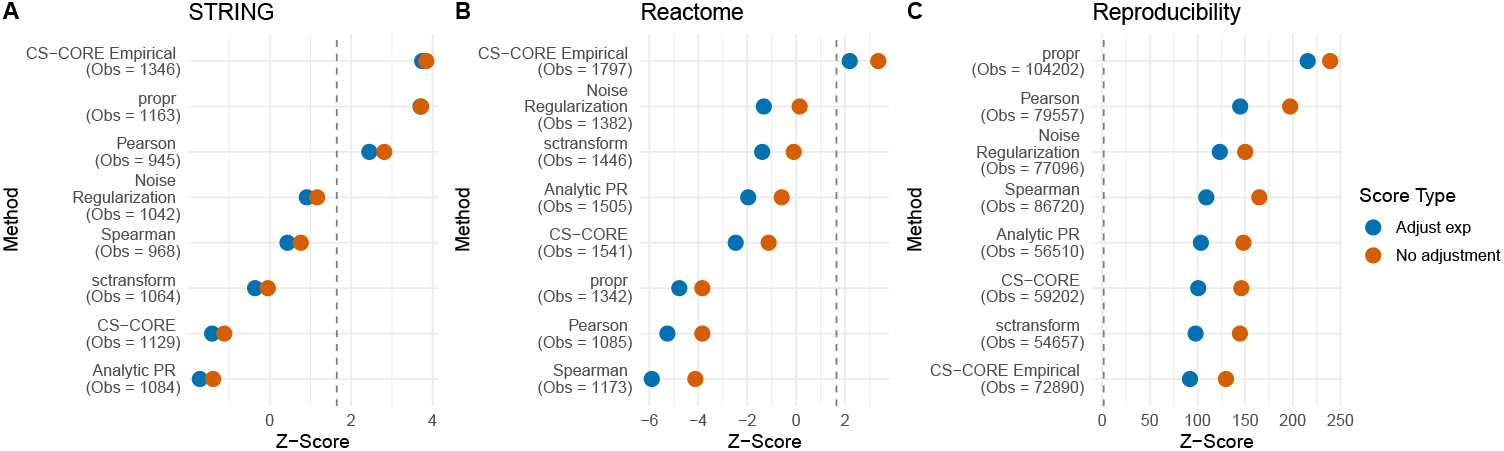
Influence of stratified adjustment on co-expression estimation method evaluation. The overlap analyses (A and B) were performed using oligodendrocytes from the ROSMAP dataset, while reproducibility (C) was assessed between oligodendrocytes from the ROSMAP and PNAS datasets. Dot plots showing the Z-scores of various co-expression estimation methods with (Adjust exp) and without (No adjustment) the stratified expression adjustment. The number of observed overlaps or reproducible pairs is shown in parentheses (Obs) for each method. The dashed line represents the significance threshold (one-sided p *<*0.05, Z-score=1.645).

**Fig. S15.**
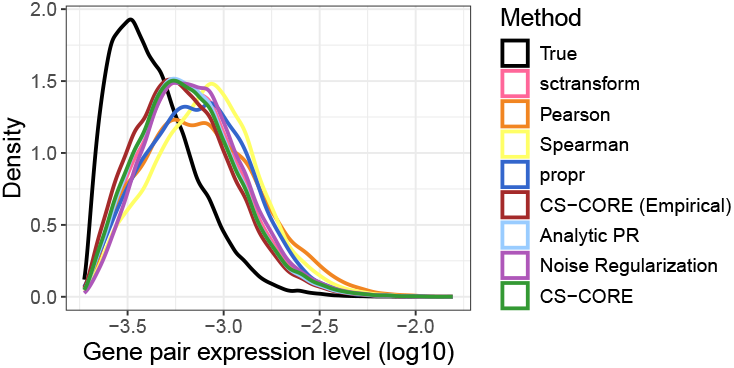
Co-expression estimation methods exhibit inherent expression-level biases. Density plots showing the distribution of mean expression levels (log10) for gene pairs identified as significant (BH adjusted p-value *<*0.05) by seven different co-expression estimation methods. For CS-CORE, we considered two different ways of calculating p-values, either through the method described in the original paper (denoted as CS-CORE) or through scSimu (denoted as CS-CORE (Empirical)). This analysis was performed on a simulated ROSMAP oligodendrocytes (see Section 4.5 for details). The distribution for the correlated pairs in the ground truth is shown in black. The plot shows that the significant pairs identified by these methods are biased towards higher mean expression levels than the true underlying distribution.

**Fig. S16.**
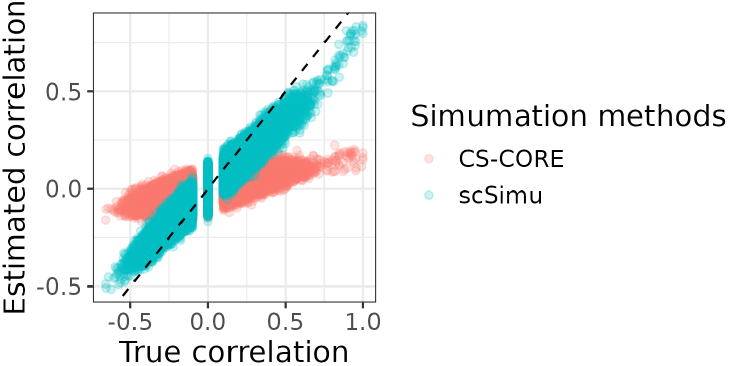
Validation of scSimu for handling non-positive semi-definite correlation matrices. We compared the ability of scSimu and the simulation method used in CS-CORE to accurately generate multivariate normal data from a non-positive semi-definite correlation matrix. The x-axis represents the true correlation values used as input, while the y-axis shows the estimated correlation from the generated multivariate normal data. The dashed line is the identity line (y=x), indicating a perfect correspondence. scSimu can better reproduce the true correlation structures, while CS-CORE simulation shows more deviations, indicating its limitation with such input matrix.

**Fig. S17.**
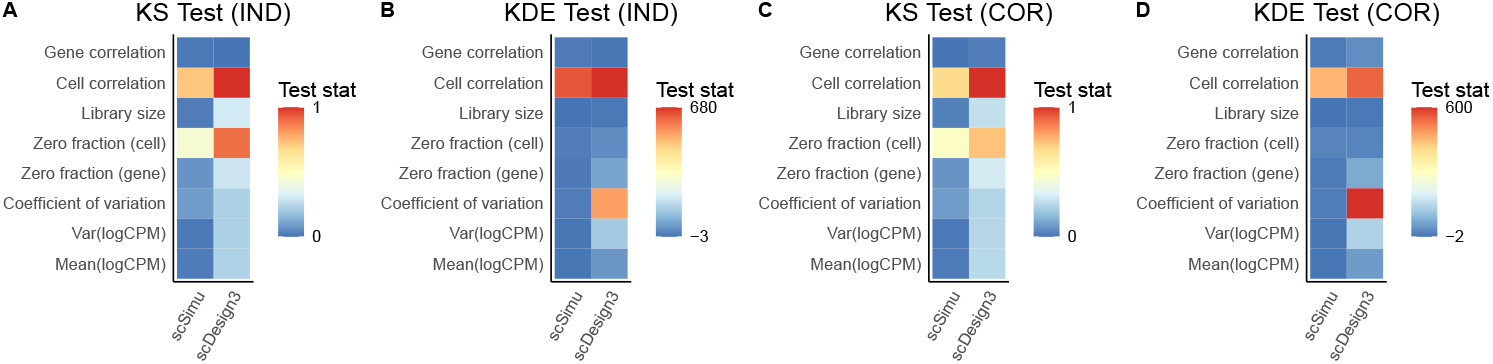
Comparison of scSimu and scDesign3 in simulating Smart-Seq data. The x-axis is the simulation method and the y-axis is the evaluation metric. Color represents the test statistics of the KS test or KDE test. The x-axis is ordered based on the mean of the test statistics. **A and B**. Evaluate the simulation of the corresponding independent data of real data based on the KS test and KDE test respectively. **C and D**. Evaluate the simulation of real data based on the KS test and KDE test respectively.

#### Non-stratified hypergeometric model

The non-stratified model quantifies the number of overlaps with the reference network when randomly selecting a subset of gene pairs. Let n be the total number of possible gene pairs, b be the number of those pairs present in a reference network, and c be the number of significant pairs identified by a method. Then, the number of overlaps by chance follows *Hypergeometric*(*n, b, c*). With the known expectation 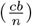 and variance 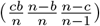, we can calculate a Z-score for the observed overlap.

The same logic can be applied to reproducibility analysis by treating the set of selected pairs from one dataset as the reference network.

#### Real data application

To assess performance against known biological networks, we applied seven different estimation methods to the top 1,000 highly expressed genes from monocytes in the lung dataset to generate gene co-expression networks. Correlated gene pairs were defined as those with a BH adjusted p-value less than 0.05. For each method, we then counted the number of overlaps with the STRING and Reactome and calculated both the stratified and non-stratified Z-scores.

We used a similar approach to assess reproducibility. We identified significant gene pairs among the top 1,000 highly expressed shared genes in oligodendrocytes from the ROSMAP and PNAS datasets. We then counted the number of significant pairs that overlap between the two cohorts and calculated both the stratified and non-stratified Z-scores to quantify reproducibility.

